# Single Turnover Transient State Kinetics Reveals Processive Protein Unfolding Catalyzed by *Escherichia coli* ClpB

**DOI:** 10.1101/2024.03.18.584833

**Authors:** Jaskamaljot Kaur Banwait, Liana Islam, Aaron L. Lucius

**Author notes:** Corresponding author Aaron L. Lucius **Email:**. **Author Contributions:** J.K.B. and A.L.L. conceived and designed experiments. J.K.B. performed experiments. J.K.B. analyzed results. J.K.B. and A.L.L wrote and revised the manuscript. J.K.B. and L.I. expressed and purified proteins. L.I. performed alphafold structure predictions of substrates. **Competing Interest Statement:** A.L.L. and J.B. are consultants for Nitrase Therapeutics.

## Abstract

*E. coli* ClpB, and *S. cerevisiae* Hsp104 are AAA+ motor proteins essential for proteome maintenance and thermal tolerance. ClpB and Hsp104 have been proposed to extract a polypeptide from an aggregate and processively translocate the chain through the axial channel of its hexameric ring structure. However, the mechanism of translocation and if this reaction is processive remains disputed. We reported that Hsp104 and ClpB are non-processive on unfolded model substrates. Others have reported that ClpB is able to processively translocate a mechanically unfolded polypeptide chain at rates over 240 amino acids (aa) per second. Here we report the development of a single turnover stopped-flow fluorescence strategy that reports on processive protein unfolding catalyzed by ClpB. We show that when translocation catalyzed by ClpB is challenged by stably folded protein structure, the motor enzymatically unfolds the substrate at a rate of ∼0.9 aa s^-1^ with a step-size of ∼60 amino acids. We reconcile the apparent controversy by defining enzyme catalyzed protein unfolding and translocation as two distinct reactions with different mechanisms of action. We propose a model where slow unfolding followed by fast translocation represents an important mechanistic feature that allows the motor to rapidly translocate up to the next folded region or rapidly dissociate if no additional fold is encountered.

## Introduction

ATPases associated with diverse cellular activities (AAA+) play essential roles in cell physiology, such as proteome maintenance, membrane fusion, DNA replication, repair and recombination, RNA processing, chromatin remodeling, and organelle biogenesis (1-5). Across domains of life, representative AAA+ molecular motors are essential for proteome maintenance (6). *E. coli* ClpB and *S. cerevisiae* Hsp104 are AAA+ protein disaggregases that, in collaboration with co-chaperones, resolve protein aggregates that form during heat shock or stress (7-12).

ClpB, in collaboration with the cochaperones, DnaK, DnaJ, and GrpE (KJE) couple the energy from ATP binding and hydrolysis to disruption of protein aggregates (7, 9, 10, 12). However, the molecular mechanisms of protein disaggregation and collaboration with cochaperones is not fully understood. ClpB is proposed to couple ATP binding and hydrolysis to processive rounds of protein unfolding and translocates the newly extracted polypeptide chain through the axial channel of the hexameric ring structure of the motor. Once passed through the axial channel, in the unfolded state, the polypeptide can refold or interact with co-chaperones that would aid in protein refolding.

Two strategies have been developed to isolate the activity of ClpB without the need for the co-chaperones, KJE. First, several “hyperactive” ClpB variants have been identified where single point mutations in the motor relieve the need for the co-chaperones, e.g. ClpB(Y503D) (13). The other strategy is using a mixture of ATP and the slowly hydrolysable ATP analogue, ATPγS. Wickner and Coworkers discovered that a 1:1 mixture of ATP:ATPγS could activate ClpB in a manner that alleviated the need to include the co-chaperones, KJE, thereby simplifying *in vitro* studies (14).

In those example *in vitro* experiments the polypeptide substrate being processed enters and leaves the reaction with structural changes but no change in molecular weight because ClpB and Hsp104 are not proteases. This fact has limited our knowledge on the mechanisms of ClpB and Hsp104 catalyzed protein unfolding and translocation. This limitation contrasts with the homologous AAA+ molecular motors, ClpA and ClpX (15, 16), which processively unfold and translocate a polypeptide into the proteolytic barrel of the associated ClpP. Thus, the product of the protein unfolding and translocation reaction catalyzed by ClpA or ClpX is covalently modified by the associated protease, ClpP. In turn, proteolytic fragments are used as a signal to report on translocation.

To interrogate ClpB and Hsp104 catalyzed polypeptide translocation we developed and applied a rapid mixing stopped-flow fluorescence method (17, 18). In that work we used unstructured polypeptides so that the time courses would reflect the kinetics of translocation and not protein unfolding. Independent of substrate length we detected only two steps before the enzymes dissociated from the unfolded polypeptides. Those results were interpreted to indicate that both ClpB and Hsp104 exhibit low processivity (P between 0.37 and 0.61) during polypeptide translocation of an unfolded chain (17, 18). However, we could not rule out the possibility that these motors might proceed through two slow steps followed by rapid translocation at a rate that is outside the millisecond temporal resolution of the stopped-flow technique. It is also possible that undetectable rapid translocation is followed by slow rate-limiting dissociation from the end.

Consistent with rapid translocation on an unfolded protein, Avellaneda *et. al*. reported translocation rates for ClpB(Y503D) to be ∼240 and 450 aa s^-1^ on mechanically unfolded substrates (19). However, neither their reported rates of ATP hydrolysis nor any published rates are consistent with this hyper-fast translocation activity (20). Similarly, Mazal *et. al*. reported ultrafast pore-loop dynamics and correlated this with rapid translocation of the unfolded κ-casein substrate (21, 22). Mazal *et. al*. acknowledged that these domain movements are substantially faster than ATP turnover.

To examine enzyme catalyzed protein unfolding, Wickner and co-workers constructed RepA(1-70)-GFP, which contains the N-terminal 70 amino acids of the phage P1 RepA protein followed by Green Fluorescent Protein (GFP) (23). The N-terminal sequence of RepA serves as a binding site for ClpB. They showed a loss of GFP fluorescence upon exposure of this construct to ClpB in the presence of a 1:1 mixture of ATP:ATPγS (14), which was interpreted to indicate ClpB unfolded GFP. They further showed no loss of fluorescence over a 45 minute time frame when only ATP or only ATPγS was provided. It is important to note that loss of GFP fluorescence does not indicate that the substrate was processively translocated after the unfolding event that led to loss of fluorescence.

Our previous examinations of ClpB and Hsp104 were carried out on unstructured polypeptide chains, so we interpret those results to reflect translocation and not protein unfolding (17, 18). Here we sought to develop a single turnover transient state kinetics approach to test for processive unfolding catalyzed by ClpB. Inspired by the RepA(1-70)-GFP constructs made by Wickner and coworkers (14, 23) and the Titin I27 substrates used extensively by the Baker and Sauer groups (24) we constructed RepA-Titin_X_, where X = 1, 2, or 3, see Fig. 1 A. In these constructs the first 70 amino acids of the RepA protein provide the binding site and are followed by one, two, or three repeats of the Titin I27 domain. Each protein ends with a C-terminal cysteine used for fluorescent labeling by maleimide chemistry.

**Figure 1:**
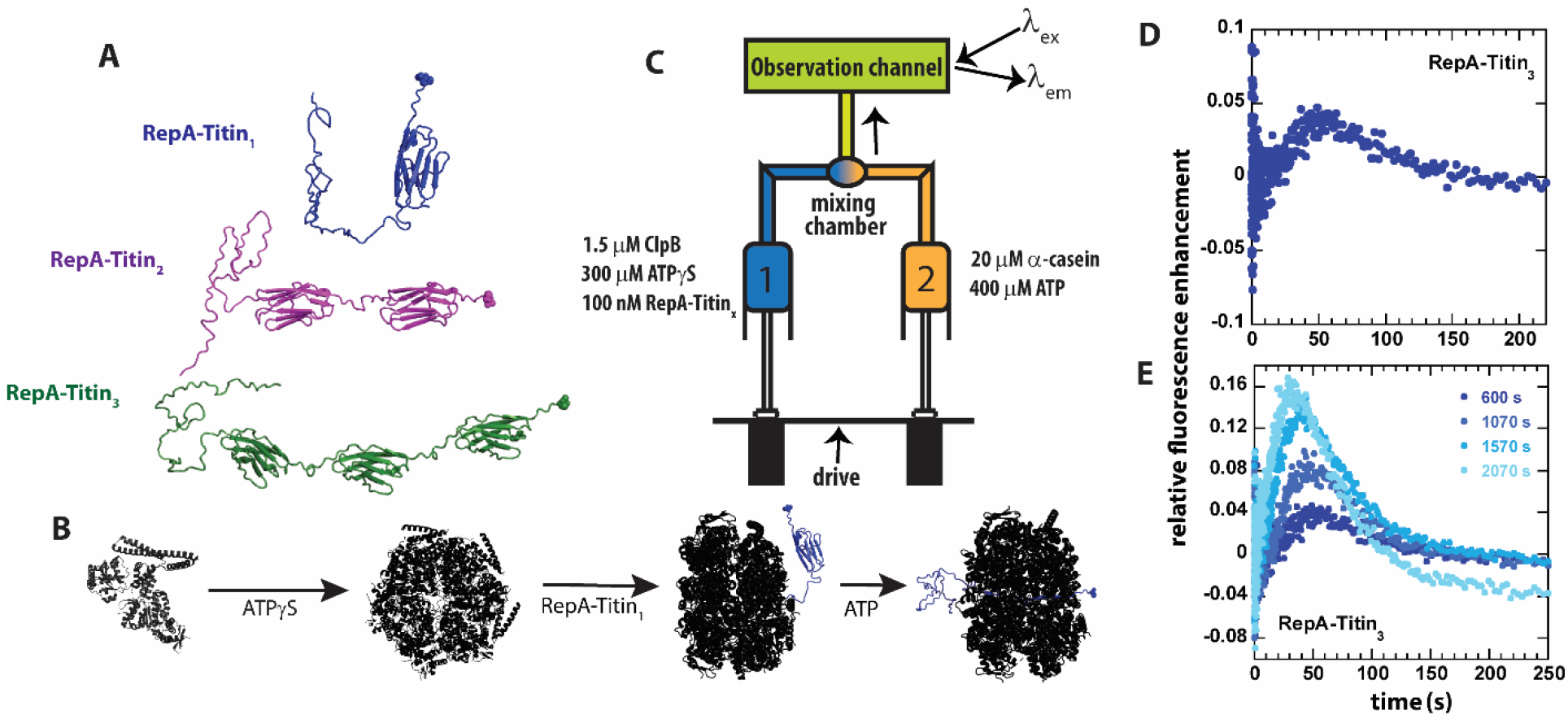
Single Turnover ClpB Catalyzed Protein unfolding. A) RepA(1-70)-Titin_1_ (Blue), RepA(1-70)-Titin_2_ (purple), RepA(1-70)-Titin_3_ (green). Each construct from N-to C-terminus consists of the first 70 amino acids of the Phage P1 RepA protein, a known binding sequence for ClpB followed by tandem repeats of the Titin I27 domain separated by linkers. Each construct contains a single cysteine shown in space-filling at the C-terminus that has been reacted with Alex Fluor (AF)-555. B) Schematic of steps in forming pre-bound complex based on our previous work (26-29). ClpB (black) is assembled into hexameric rings competent for substrate binding by adding ATPγS, illustrated as bound to the RepA-Titin_1_ substrate in blue, followed by rapid mixing with ATP. As shown, ClpB is expected to unfold the Titin I27 domains and translocate the newly unfolded substrates through the axial channel of the hexameric ring. C) Schematic representation of stopped-flow. Syringe 1 contains the indicated concentrations of ClpB monomer, ATPγS, and RepA-Titin_X_, where x = 1, 2, 3. Syringe 2 contains 400 μM ATP and 20 μM α-casein to serve as a trap for any free ClpB. The contents of the two syringes are rapidly mixed at a 1:1 mixing ratio and flow into the observation channel where AF555 is excited at λ_ex_ = 555 nm and emission is observed at λ_em_ >570 nm. D) Representative time-course collected using strategy in (C) using RepA-Titin_3_ after pre-incubating the sample at 25 °C for 600 s. E) Successive experimental time-courses collected as in D. The total time of incubation before collection of the time-course is indicated.

Using the RepA-Titin_X_ constructs, we report evidence of sequential and processive unfolding of the tandem repeats of the stably folded Titin I27 domains. Surprisingly, we have also found that ATPγS alone will support processive protein unfolding and translocation of these constructs. Because of this we developed a sequential mixing stopped-flow strategy to separate and quantify the rates of ATPγS and ATP:ATPγS driven protein unfolding. Here we report rates of protein unfolding in the range of 1 – 4 amino acids (aa) s^-1^ and ∼60 aa are unfolded during each rate limiting unfolding event. These rates of protein unfolding are approximately two orders of magnitude slower than the reported translocation rates on unfolded polypeptide chains reported by others (19, 22). Thus, we propose that protein unfolding catalyzed by ClpB is rate-limiting and, upon unfolding, translocation on the newly unfolded polypeptides is much faster than protein unfolding. Our method reveals mechanistic insights into ClpB catalyzed protein unfolding that have been inaccessible by other techniques. Importantly, our technology overcomes the barrier of needing covalent modification to detect enzyme catalyzed protein unfolding and translocation by the protein disaggregating machines. The approach can be broadly applied to the many AAA+ motors that have been hypothesized to catalyze protein unfolding and translocation but do not covalently modify the substrate on which they operate.

## Results

### Development of Single Turnover Protein Unfolding Method

To test for protein unfolding catalyzed by wild type (wt) ClpB, we engineered constructs containing the N-terminal 70 amino acids of the RepA protein (14, 25) followed by tandem repeats of the Titin I27 domain. Each construct contains a single cysteine residue at the C-terminus labeled with Alexa Fluor (AF) 555-maleimide, see Fig. 1 A. The rationale for these constructs is that we can preassemble ClpB into the biologically active hexamers in the presence of ATPγS (26-28) and bind the hexamer to the N-terminal RepA sequence, (29) see Fig 1 B for schematic of assembly and binding steps.

The pre-bound complex is loaded into Syringe 1 of the stopped-flow fluorometer. In the other syringe is ATP and excess α-casein. The α-casein serves as a trap for any unbound ClpB, see Fig. 1 C for schematic of stopped-flow setup. The contents of the two syringes are rapidly mixed within 2 ms, AF555 is excited at λ_ex_ = 555 nm, and emission is observed at λ_em_ > 570 nm.

After mixing, if ClpB processively unfolds the Titin I27 domain and translocates N to C, then, upon arrival of ClpB at the AF555, we anticipate protein-induced fluorescence enhancement (PIFE) (30), which has been reported to occur with AF555 (31). In addition, PIFE was tested using an unstructured 50 aa polypeptide labeled with AF555. When ClpB binds this unstructured 50-mer, PIFE is observed due to the close proximity of AF555 to ClpB, see Supp. Fig 1 (17, 29).

Experiments were performed by stopped-flow mixing of 1.5 μM ClpB, 300 μM ATPγS, and 100 nM RepA-Titin_3_ with 20 μM α-casein and 400 μM ATP, see Fig. 1 C for mixing schematic. Upon mixing, fluorescence emission from AF555 was monitored as a function of time, see Fig. 1 D. The time-course exhibits a lag phase followed by fluorescence enhancement and gradual loss of signal. The lag is interpreted to indicate the time over which ClpB is unfolding the Titin I27 domains and translocating the newly unfolded polypeptide chain. The fluorescence enhancement is interpreted to indicate arrival of the motor at the C-terminus and induction of PIFE followed by loss of signal due to dissociation.

In these stopped-flow experiments, one fills the contents of the two syringes illustrated in Fig. 1 C and collects multiple time-courses using the same sample upon successive mixing events. The only difference from mixing event to mixing event is that the sample will have aged for the time it took to collect the previous time-course/courses. Thus, the time-courses are expected to overlay between mixing events if the contents of the two syringes are at chemical and thermal equilibrium. Thermal equilibrium is achieved by maintaining contact with a temperature-controlled water bath set to 25 °C and sufficient incubation time. Chemical equilibrium is judged by comparing time-courses collected in succession.

Here we observed systematic deviation in the collected time-courses from mixing event to mixing event, see Fig. 1 E and a zoomed-in version of Fig 1 E shown in Supp. Fig 2. The first collected time-course (dark blue) is labeled 600 s as the sample was first allowed to preincubate for 10 minutes to achieve thermal equilibrium before collection, see Fig. 1 D and E. Select, subsequent time-courses are shown, and the total pre-incubation time is indicated, see Fig. 1 E and Supp. Fig. 2. For each successive collection the lag time is getting shorter, the emergence of a peak occurs earlier in time, and the magnitude of the peak is increasing. The latter is most consistent with binding equilibrium not being fulling achieved. But the former two observations suggest that the longer the samples are incubated the less time it takes before PIFE after mixing with ATP. This phenomenon was also observed with RepA-Titin_1_ and RepA-Titin_2_, data not shown. These observations suggest that ClpB is moving closer to the fluorophore during the pre-incubation time.

**Figure 2:**
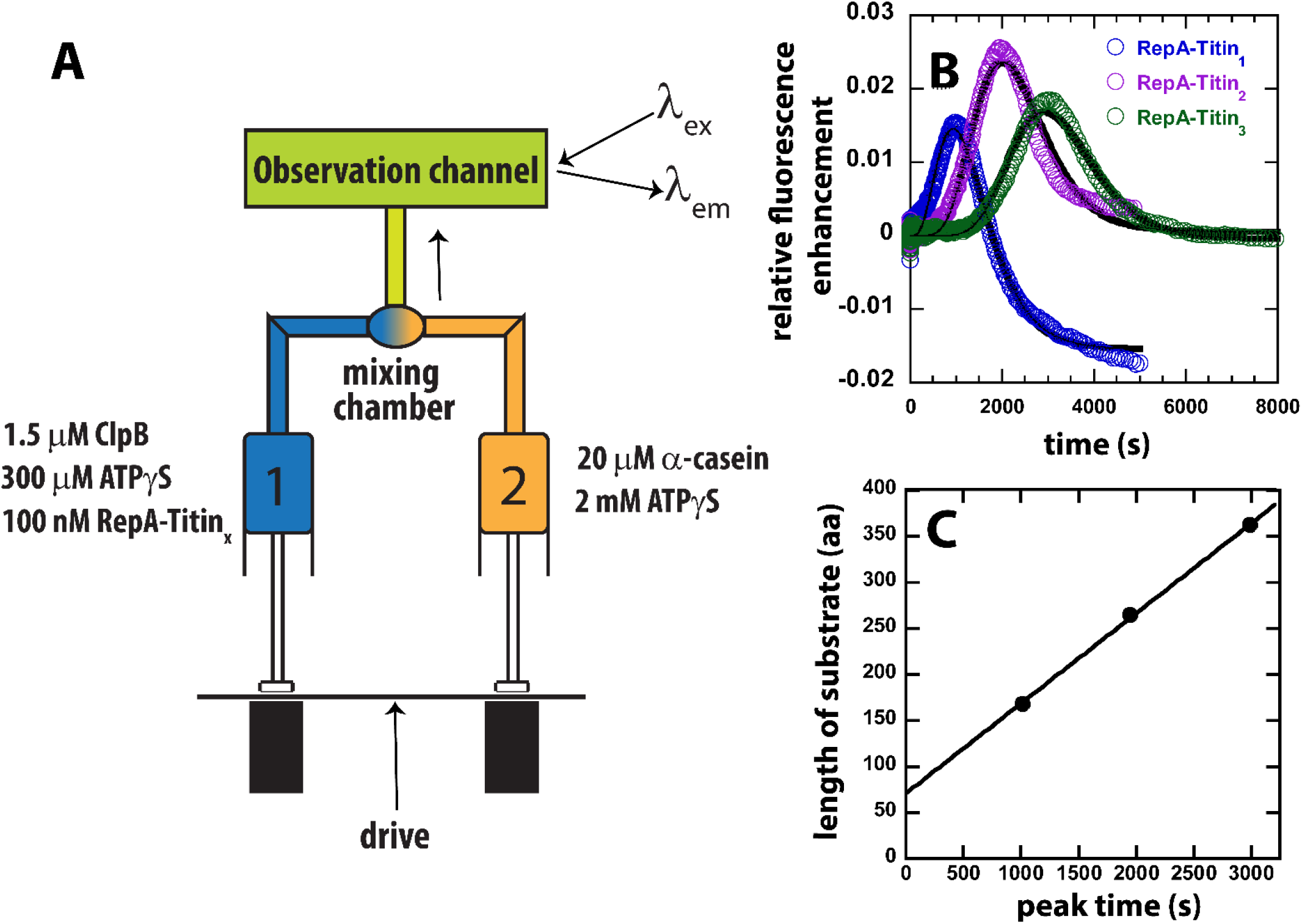
Test for ATPγS driven Protein Unfolding by ClpB. A) Mixing strategy as in Fig. 1 C but with ATP replaced with 2 mM ATPγS in Syringe 2. B) Time-courses collected using RepA-Titin_1_(blue), RepA-Titin_2_(purple), and RepA-Titin_3_(green) plotted as relative fluorescence enhancement vs. time. The solid black line represents the best-fit line from fitting to Scheme 1, Fig. 4 A. The fitting parameters obtained are the unfolding rate constant, *k*_*U*_ = (0.0042 ± 0.0003) s^-1^, and the kinetic step-size, *m* = 26 (21, 28) aa. C) length of substrate vs. peak time determined from B. The plot was fit to a linear equation to yield a slope = (0.098 ± 0.003) aa s^-1^ and intercept of (71 ± 7) aa.

We hypothesized that the variability observed in Fig. 1 E and Supp. Fig. 2 was due to slow ATPγS hydrolysis being coupled to protein unfolding and translocation catalyzed by ClpB. Consequently, during the time between mixing events some pre-unfolding and pre-translocation from the N-terminal binding site to the C-terminal fluorophore had already occurred in the syringe.

To test for ATPγS driven unfolding and translocation, we loaded ClpB pre-bound to RepA-Titin_1_ in the presence of 300 μM ATPγS into Syringe 1 of the stopped-flow. In Syringe 2, we loaded 2 mM ATPγS and 20 μM α-casein trap with no hydrolysable ATP, see Fig. 2 A. The rationale for this design is that, even though we anticipate pre-translocation due to the presence of 300 μM ATPγS, we expect translocation to be accelerated upon mixing and increasing the total concentration of ATPγS.

In the absence of hydrolysable ATP but in the presence of only ATPγS, the time-courses exhibit a distinct lag, followed by a fluorescence enhancement, and a slow loss in signal, see Fig. 2 B. Consistent with translocation from the N-terminal RepA sequence to the C-terminal fluorophore both the lag time and the time of appearance of the peak is observed to increase for each addition of another Titin I27 domain, see Fig. 2 B. This indicates that ClpB proceeds through an increasing number of rate-limiting steps for each increase in substrate length.

If the observed peak in the time-courses for each RepA-Titin_X_ substrate, in Fig. 2 B, represent arrival of ClpB at the C-terminal fluorophore then plotting the total length of the substrate vs. emergence of the peak, termed “peak time”, should represent a classical kinematics position vs. time plot. Fig. 2 C shows the length of the substrate vs. the peak time from the time-courses in Fig. 2 B. As expected, substrate length vs. peak time exhibits a linear increase with a slope of (0.098 ± 0.003) aa s^-1^ and an intercept of (71 ± 7) aa. From these observations we hypothesize that the slope represents the rate of unfolding and translocation in the presence of a final mixing concentration of 1.15 mM ATPγS and no ATP. We propose that the intercept represents the number of amino acids pre-unfolded and pre-translocated in the presence of 300 μM ATPγS during the ten minute pre-incubation time.

To further test ATPγS driven translocation, we performed a series of experiments where only ClpB was loaded into Syringe 1 and, in Syringe 2 was loaded ATPγS and RepA-Titin_X_, see Supp. Fig. 3 A for mixing schematic. In this setup, before PIFE can be observed, ClpB must bind ATPγS, assemble into hexamers, bind to the RepA sequence, and initiate protein unfolding and translocation. Importantly, these experiments may not be single turnover because the ClpB is not pre-bound to the substrate and there is no trap for free ClpB, i.e. no α-casein. Nevertheless, the time-courses do exhibit a lag phase that increases with increasing substrate length, followed by PIFE, and an apparent plateau, see Supp. Fig. 3 B. As expected, when assembly and binding must occur before protein unfolding and translocation can ensue the time-courses exhibit a longer lag and undetectable dissociation after arrival at the fluorophore, compare time courses in Supp. Fig. 3 B to Fig. 2 B. In sum, all experiments in the presence of only ATPγS as an energy source reveal that ClpB processively unfolds tandem repeats of Titin I27.

**Figure 3:**
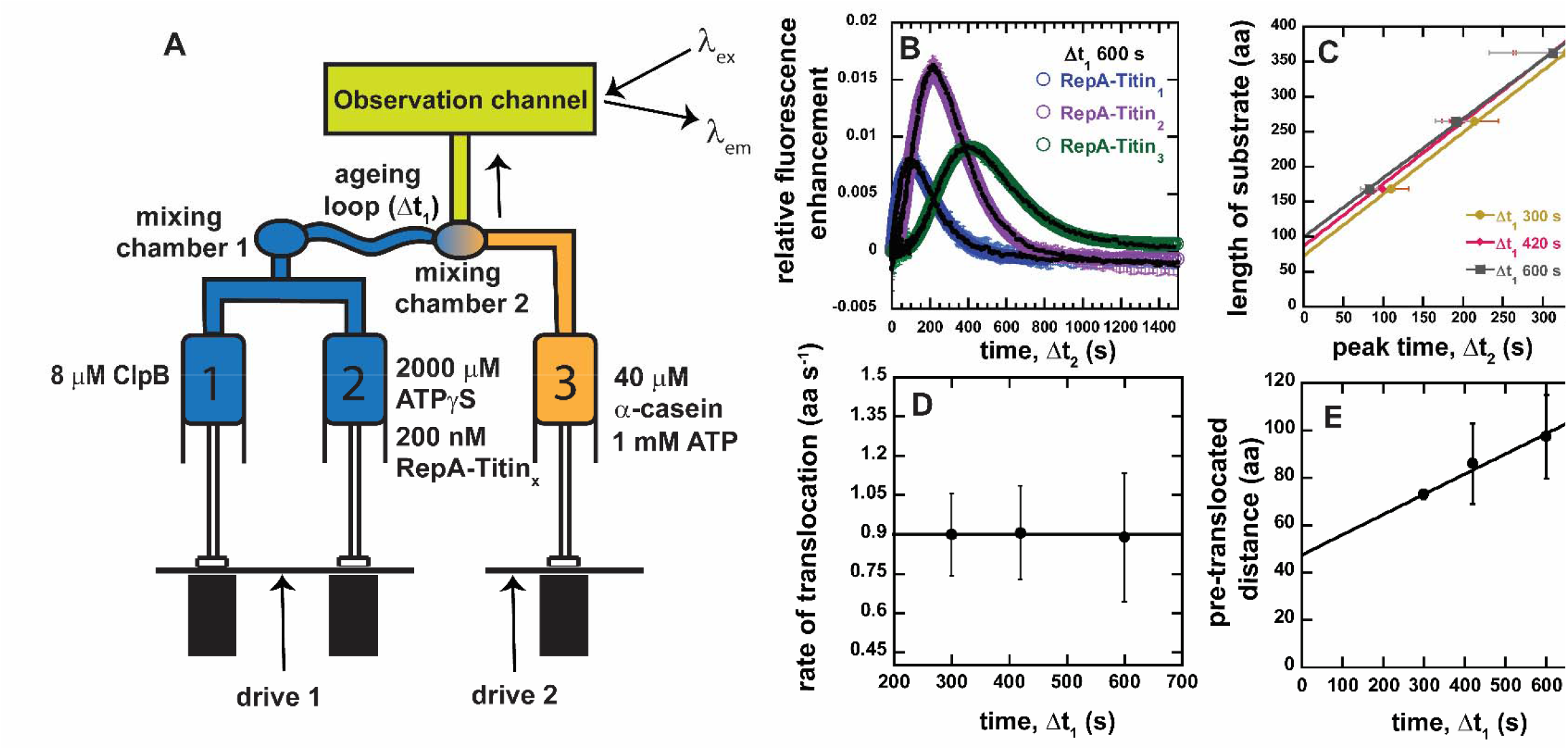
Sequential Mixing Stopped-Flow Strategy. A) Schematic representation of sequential mixing. Syringe 1 contains 8 μM ClpB monomer, Syringe 2 contains 2 mM ATPγS and 200 nM RepA-Titin_X_, Syringe 3 contains 40 μM α-casein and 1 mM ATP. The contents of Syringes 1 and 2 are mixed 1:1 in mixing chamber 1 leading to a concentration of 4 μM ClpB, 1 mM ATPγS, and 100 nM RepA-Titin_X_. The sample ages for a user-defined amount of time, Δt_1_, followed by rapid mixing with the contents of Syringe 3. The sample flows into the observation channel at a final concentration of 2 μM ClpB, 50 nM RepA-Titin_X_, 500 μM ATPγS, 500 μM ATP, and 20 μM α-casein. In the observation channel AF555 is excited at λ_ex_ = 555 nm and emission is observed at λ_em_ >570 nm. B) Representative time-courses from the average of five or more sequentially collected time-courses for RepA-Titin_1_(blue), RepA-Titin_2_(purple), and RepA-Titin_3_(green) at Δt_1_ = 600 s. The black solid lines represent the best-fit line from fitting to Scheme 1 (see Fig. 4 A). The fitting parameters obtained are *k*_*U*_ = (0.017 ± 0.002) s^-1^ and *m* = (56.5 ± 0.7) aa. C) Total length of substrate as a function of peak time determined for Δt_1_ = 300, 420, and 600 s. Solid lines represent weighted linear fits yielding D) slope vs. Δt_1_ and E) the intercept vs. Δt_1_. All data points and error bars represent the average and standard deviation determined from three replicates.

### Development of Sequential Mixing Strategy to Control and Quantify ATPγS Driven Protein Unfolding and Translocation

ClpB is pre-bound to the polypeptide chain to remove the kinetics of ClpB assembly and binding to the protein substrate. This is important as we are seeking to acquire time-courses that are rate-limited only by the kinetics of protein unfolding and/or translocation but not the kinetics of hexamer formation or substrate binding.

Previously we reported that only ATPγS would support ClpB binding to the polypeptide and the non-hydrolysable nucleoside triphosphate analogues, AMPPNP and AMPPCP could not substitute for ATPγS (28). Consequently, we cannot eliminate the ATPγS pre-unfolding/pre-translocation if we seek to have a pre-bound complex. Thus, we devised a sequential mixing strategy to account for and control the pre-unfolding and translocation that occurs in the presence of ATPγS.

Sequential mixing stopped-flow allows for two mixing events, illustrated in Fig. 3 A. In this design we load 8 μM ClpB monomer into Syringe 1 and, in Syringe 2, we load 2 mM ATPγS and 200 nM RepA-Titin_X_, see Fig. 3 A for mixing schematic. The contents of Syringes 1 and 2 are rapidly mixed and allowed to age, without observation, for a user-defined amount of time, Δt_1_. In this experiment, the Δt_1_ is the amount of time where ClpB will bind ATPγS, assemble into hexamers, bind to RepA-Titin_X_ and initiate some pre-unfolding and pre-translocation.

After Δt_1_ elapses the sample is rapidly mixed with the contents of the third syringe containing 40 μM α-casein and 1 mM ATP, see Fig. 3 A. After the second mixing event the sample flows into the observation chamber and fluorescence as a function of time, Δt_2_, is monitored. The final concentrations after the two mixing events are 2 μM ClpB monomer, 50 nM RepA-Titin_X_, 500 μM ATPγS, and 500 μM ATP. The final ratio of ATP to ATPγS was chosen because it has been shown that ClpB exhibits its highest protein disaggregation activity and protein unfolding activity in the presence of a 1:1 mix of ATP:ATPγS. Moreover, the 1:1 mix activates ClpB in the absence of the co-chaperones, KJE (14).

A series of representative time-courses collected with Δt_1_ = 600 s are shown in Fig. 3 B. As expected, we observe a lag and a maximum fluorescence that increases in time with increasing number of Titin I27 domains. Importantly, the representative time-courses in Fig. 3 B for each RepA-Titin_X_ represent the average of five or more successive time-courses collected using the same sample preparation. Thus, in this design, no variability between mixing events is detected since ClpB and ATPγS has been pre-incubated for precisely the same amount of time, Δt_1_, for each successive mixing event, see Supp. Fig. 4.

**Figure 4:**
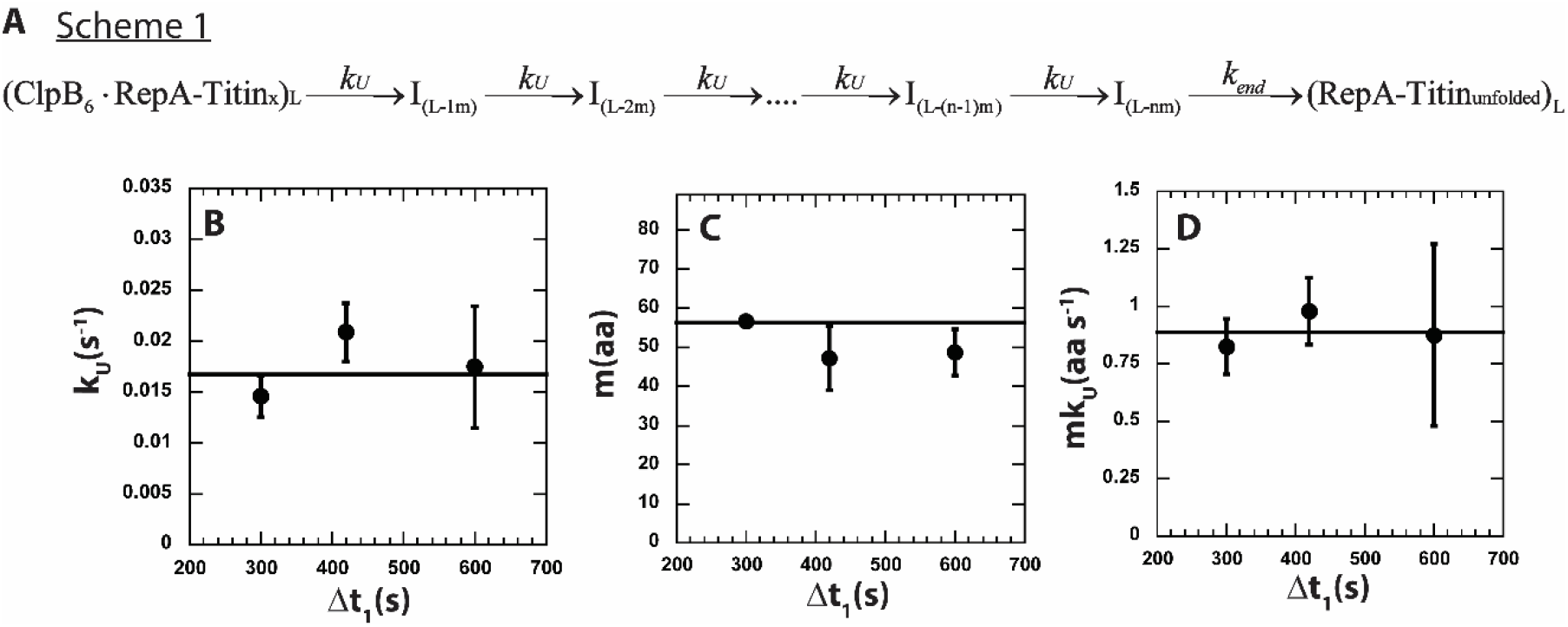
*n*-step sequential mechanism for 500:500 μM ATP:ATPγS. A) Proposed kinetic scheme for ClpB catalyzed protein unfolding and translocation on RepA-Titin_X_ substrates. Parameters B) *k*_*U*_ C) *m* D) *mk*_*U*_ obtained from fitting to scheme 1 at each Δt_1_ are shown in solid black circles. The black solid line represents the best weighted fit line to a linear equation with zero slope to yield the average value of the unfolding rate constant, *k*_*U*_ = (0.017 ± 0.002) s^-1^, kinetic step-size, *m* = (56.5 ± 0.7) aa, and overall rate, *mk*_*U*_ = (0.89 ± 0.09) aa s^-1^.

In all cases, the time courses collected with RepA-Titin_2_ exhibit higher peak max values compared to RepA-Titin_1_ and RepA-Titin_3_. This is because all three substrates are fluorescently labeled to a different labeling efficiency and thus there is a different proportion of labeled to unlabeled substrate in each sample. ClpB will bind to both the labeled and unlabeled substrates and the presence of unlabeled substrate will compete for binding with labeled substrate. RepA-Titin_2_ exhibits the highest labeling efficiency and thus the lowest concentration of unlabeled substrate to compete for binding. Thus, time courses collected with RepA-Titin2 always exhibits the highest peak max since the peak max is proportional to extent of binding, see discussion below.

In this sequential mixing design, ClpB will still pre-unfold and pre-translocate the substrate during Δt_1_. To further control and account for ATPγS driven translocation, we varied Δt_1_ to be 300, 420, and 600 s. The substrate length vs. peak time for each Δt_1_ is plotted and fit to a line, see Fig 3 C. The slope and intercept for each value of Δt_1_ is plotted in Fig. 3 D and E, respectively.

Fig. 3 D indicates that the rate is independent of Δt_1_ within the time range tested. The average value from the plot in Fig. 3 D is (0.9 ± 0.1) aa s^-1^ and is taken as the rate of translocation in the presence of a 1:1 mix of ATP to ATPγS, in this case 500 μM:500 μM.

The intercept values from the fits in Fig. 3 C increase linearly with increasing Δt_1_, see Fig. 3 E. We hypothesize that the intercept represents the number of amino acids pre-unfolded and pre-translocated in the presence of 1 mM ATPγS during Δt_1_, which we term pre-translocated distance. The slope of the pre-translocated distance vs. Δt_1_ yields a value of (0.09 ± 0.06) aa s^-1^, consistent with the rate of translocation determined in only the presence of 1.15 mM ATPγS in Fig. 2. Interestingly, the weighted linear fit in Fig. 3 E yields an intercept of (48 ± 17) amino acids when extrapolating to Δt_1_ = 0, see Table 1. We interpret this number to represent the average number of amino acids that are not involved in unfolding and translocation, which we will term the excluded length. The excluded length may represent the number of amino acids that are in contact with the motor but not part of the lattice to be translocated, the number of amino acids dangling outside of the hexameric ring relative to the direction of translocation, or some combination of the two, see Supp. Fig. 5 for schematic representation of these various distances.

**Table 1.** Parameters obtained from model-independent analysis on experiments presented in Fig. 3 and Supp. Fig. 7.

| Parameters from model-independent analysis | 500 $\mu\text{M}$ [ATPγS] and 500 $\mu\text{M}$ [ATP] | 150 $\mu\text{M}$ [ATPγS] and 500 $\mu\text{M}$ [ATP] |
| --- | --- | --- |
| Rate of translocation with ATPγS and ATP (aa s$^{-1}$) | $0.9 \pm 0.1$ | $4.3 \pm 0.2$ |
| Excluded length (aa) | $48 \pm 17$ | $67 \pm 23$ |
| Rate of translocation with ATPγS (aa s$^{-1}$) | $0.09 \pm 0.06$ | $0.05 \pm 0.05$ |

**Figure 5.**
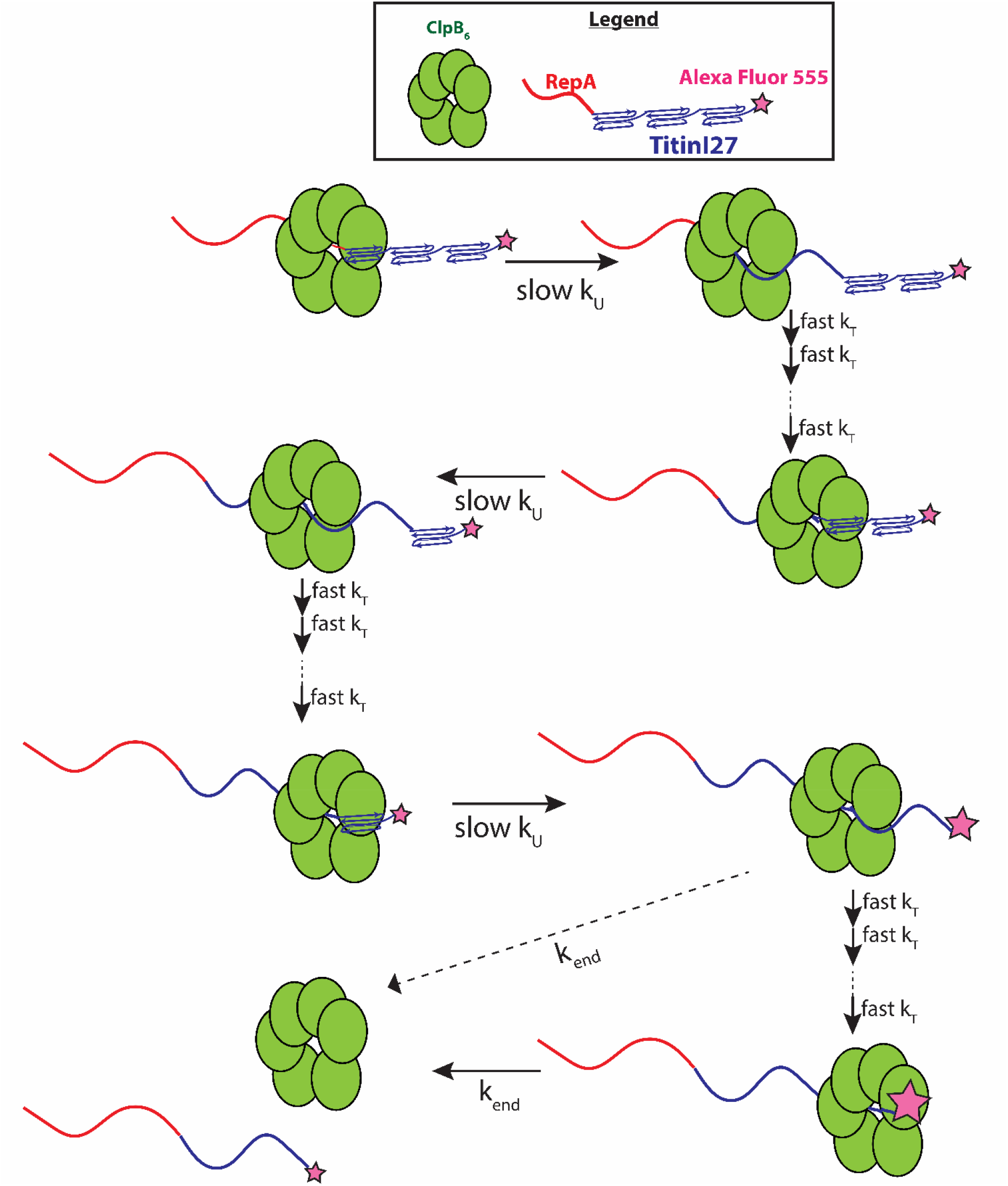
Proposed mechanism of ClpB catalyzed protein unfolding and translocation. In the presence of ATPγS, hexameric ClpB binds to the unfolded 70 amino acid long RepA sequence. Upon mixing with ATP, ClpB unfolds ∼60 amino acids with rate constant, k_U_, shown as complete collapse of the first Titin I27 domain. ClpB then proceeds through multiple fast translocation steps with rate constant k_T_ before arrival at the next folded Titin I27 domain and the process repeats. After complete unfolding of the final Titin I27 domain, ClpB may dissociate before arrival at the C-terminus, or rapidly translocate to the C-terminus followed by slow dissociation with dissociation rate constant k_end_. We expect PIFE to occur the last unfolding step and/or at the last translocation step; the relative intensities of AF555 represented by the size of the pink star.

### Global Fitting of Length-Dependent Time-courses

Scheme 1, in Fig. 4 A, represents an *n*-step sequential mechanism that we propose to describe the experimental time-courses collected at each value of Δt_1_, see Fig. 3 B for Δt_1_ = 600 s. Scheme 1 (see Fig. 4 A) shows ClpB pre-bound to a RepA-Titin_X_ of substrate length, *L*. Upon mixing with ATP the motor proceeds through *n* number of unfolding steps with rate constant *k*_*U*_. Each intermediate is denoted as *I*_*(L-im)*_, where *L* represents the total number of amino acids in the substrate, *i* is the number of steps taken to arrive at a given intermediate, *I*, and *m* represents the average number of amino acids unfolded between two rate limiting steps, i.e. the kinetic step-size. The rate constant, *k*_*end*_, represents dissociation from the C-terminal end of the unfolded polypeptide chain denoted as (RepA-Titin_unfolded_)_L_.

The observed total number of steps, *n*, is hypothesized to be proportional to the substrate length divided by the kinetic step-size as *n = L/m*. To test this, the three time-courses were simultaneously fit using Scheme 1 (see Fig. 4 A and Equation 4) with *k*_*U*_ constrained to be the same for all three time-courses, i.e. global, and the number of steps, *n*, and *k*_*end*_ local to each time-course. In this analysis, we ascribe signal change to the last intermediate, *I*_*(L-nm)*_, and the unfolded product, (RepA-Titin_unfolded_), see Materials and Methods.

The determined number of steps as a function of substrate length is plotted in Supp. Fig. 6 A. The n vs. L plot was fit to a line with a slope of (0.016 ± 0.002) steps aa^-1^ and an intercept of (−1.48 ± 0.42) steps. As seen in Supp. Fig 6 A, the line of best fit exhibits an x-intercept of ∼93 amino acids. We interpret the intercept to indicate that an average of ∼93 amino acids are pre-unfolded during Δt_1_ = 600 s. Consistently, the plot of pre-translocated distance vs. Δt_1_, in Fig. 3 E, shows that the average of the pre-translocated distance across three replicates collected with Δt_1_ = 600 s is ∼97 amino acids. This result suggests that ∼97 amino acids are pre-unfolded or pre-translocated in the presence of ATPγS before mixing with ATP.

The simple relationship between the number of steps, *n*, and the substrate length, *L*, is *n = L/m*. However, due to pre-translocated distance during Δt_1_, some of the total length of the substrate has already been translocated. To account for this, we define the reduced length, *L’* = (*L* – *C*), where *C* represents the number of amino acids pre-translocated for a given Δt_1_. Thus, the number of steps, *n*, is related to reduced length as *n* = (*L* – *C*)/*m* = *L’*/*m*, see Supp. Fig. 5 for schematic.

Thus, for all of the length-dependent time-courses collected with variable Δt_1_ the number of steps, *n* in Eq. 5 was replaced with (*L* – *C*)/*m*, where *C* was determined independently for each replicate, see Materials and Methods. The time-courses were simultaneously fit with *m* and *k*_*U*_ as global parameters and *k*_*end*_ as a local parameter.

The rate constant describing dissociation, *k*_*end*_, was local to each time-course because we observed local variability in this phase. This is not surprising as this phase represents dissociation and fluorescence changes due to refolding. Although the impact on the signal due to dissociation is likely the same for each substrate, the influence of refolding on the signal is probably not the same because of the differences in total length.

The solid lines in Fig. 3 B represent the best fit lines and indicate excellent agreement between the model and the experimental time-courses. From the global fit of each of three replicates for three Δt_1_ values we found the unfolding rate constant, *k*_*U*_, the kinetic step-size, *m*, and the overall rate, *mk*_*U*_ to all be independent of Δt_1_, see Fig. 4 B -D. As a function of Δt_1_ we determined an average kinetic step-size of m = (56.5 ± 0.7) aa step^-1^ indicating that ∼57 amino acids are unfolded between two rate limiting steps with unfolding rate constant, *k*_*U*_ = (0.017 ± 0.002) s^-1^, see Table 2. The product of *m* and *k*_*U*_ yields the overall rate of *mk*_*U*_ = (0.89 ± 0.09) aa s^-1^, which is in excellent agreement with (0.9 ± 0.1) aa s^-1^ determined from the peak time analysis in Fig. 3 D.

**Table 2.** Parameters obtained from global fitting the time-courses obtained from experiments in Fig. 3 and Supp. Fig. 7 to Scheme 1 in Fig. 4 A.

| Parameters from model-dependent analysis | 500 $\mu\text{M}$ [ATPγS] and 500 $\mu\text{M}$ [ATP] | 150 $\mu\text{M}$ [ATPγS] and 500 $\mu\text{M}$ [ATP] |
| --- | --- | --- |
| $m(\text{aa})$ | $56.5 \pm 0.7$ | $58 \pm 4$ |
| $k_U(\text{s}^{-1})$ | $0.017 \pm 0.002$ | $0.055 \pm 0.005$ |
| $mk_U(\text{aa s}^{-1})$ | $0.89 \pm 0.09$ | $3.1 \pm 0.2$ |

### Impact of the ATP:ATPγS Ratio on Processive Unfolding and Translocation

To test the impact of different ATP:ATPγS mixing ratios, experiments were carried out as schematized in Fig. 3 A but with a lower concentration of ATPγS. As in Fig. 3 A, 8 μM ClpB was loaded into Syringe 1 but 600 μM ATPγS and 200 nM RepA Titin_X_ substrate were loaded into Syringe 2. Syringe 3, again, contains 1 mM ATP and 40 μM α-casein. Thus, after the two mixing events, the final concentration of ATP and ATPγS were 500 μM and 150 μM, respectively, yielding an approximately 3:1 ATP:ATPγS mixing ratio.

Supp. Fig. 7 A shows a series of representative time-courses for all three RepA-Titin_x_ substrates at fixed Δt_1_ = 600 s. The relative fluorescence enhancement value is approximately two-fold lower and the time-courses exhibit larger fluctuations in the signal compared to time-courses collected at 1:1 ATP:ATPγS, compare Supp. Fig. 7 A to Fig 3 B.

The amplitude of a single turnover experiment is directly proportional to the amount of bound enzyme or the extent of binding, 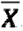, given by Eq. 1 (32), where **[*ClpB***_**6**_ ***-RepA -Tilin***_***x***_] represents hexameric ClpB bound to the RepA-Titin_X_ substrate and **[*RepA – Tilin***_***x***_]_***Total***_ represents the total amount of RepA-Titin_X_ present in the experiment. In this single-turnover experiment, the peak height is expected to be proportional to the amount of ClpB that arrived at the fluorophore after mixing with ATP. However, the amount of ClpB that made it to the fluorophore is defined by the processivity, *P*, given by Eq. 2, where *k*_*U*_ is as defined in Scheme 1 (see Fig. 4 A) and *k*_*d*_ is the rate constant for dissociation at each intermediate (32). Thus, the peak height is proportional to the amount of ClpB initially bound,, times the processivity, *P*, or, given in Eqs. 1 and 2

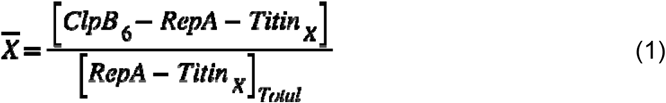

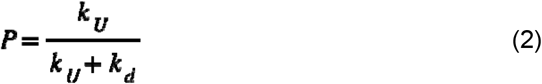

The observed reduction in peak height upon reducing the [ATPγS] is consistent with less ClpB bound, lower processivity, or both. Less ClpB bound is expected as we have shown that the fraction of hexameric ClpB present in solution is a function of both ATPγS concentration and ClpB concentration (26, 27). Thus, less hexamer is present and able to bind the substrate at 150 μM ATPγS compared to 500 µM ATPγS. However, reduced processivity at lower ATPγS may also be contributing to the lower amplitude.

To determine the rate of ClpB catalyzed protein unfolding in the presence of approximately 3:1 ATP:ATPγS we repeated the experiment at different values of Δt_1_, plotted substrate length vs. peak time, and performed the same peak time analysis as in Fig. 3, see Supp. Fig. 7 B – D.

Interestingly, the rate of translocation in the presence of approximately 3:1 ATP:ATPγS is (4.3 ± 0.2) aa s^-1^, see Supp. Fig. 7 C and Table 1. This is ∼ 5 times faster than the ∼0.9 aa s^-1^ in the presence of a 1:1 ATP:ATPγS, see Table 1. This suggests that although, on the one hand, ATPγS activates ClpB, ATPγS is also competing with ATP and slowing unfolding and/or translocation.

Supp. Fig. 7 D shows the intercept values (pre-translocated distance) from the fits in Supp. Fig. 7 B plotted as a function of Δt_1_. The intercept, pre-translocated distance, increases with increasing time, Δt_1_, again consistent with ATPγS driven pre-unfolding/translocation. The rate is ∼0.05 aa s^-1^, which represents the rate of translocation in the presence of 150 μM ATPγS. Consistently, this is slower than the ∼0.09 aa s^-1^ in the presence of 500 μM ATPγS although the values are within error. Interestingly, the intercept in Supp. Fig. 7 D is ∼67 amino acids in the presence of 150 μM ATPγS, which is larger than the ∼48 amino acids in the presence of 500 μM ATPγS. However, as can be seen in Table 1, this number has large uncertainty, presumably, due to the long extrapolation back to Δt_1_ = 0.

The time-courses in Supp. Fig. 7 A were also subjected to global analysis using Scheme 1 in Fig. 4 A. The resultant kinetic parameters as a function of Δt_1_ are shown in Supp. Fig. 8 A – C. From this analysis we determined a kinetic step-size of *m* = (58 ± 4) aa step^-1^ indicating that at these lower ATPγS concentrations we detect ∼58 amino acids to be unfolded between two rate limiting steps with unfolding rate constant, *k*_*U*_ = (0.055 ± 0.005) s^-1^. The overall rate of unfolding, *mk*_*U*_ = (3.1 ± 0.2) aa s^-1^, which is in reasonable agreement with (4.3 ± 0.2) aa s^-1^ determined from the peak time analysis, see Tables 1 and 2.

## Discussion

Substantial evidence exists indicating that ClpA processively unfolds and translocates a polypeptide through its axial channel, out the other side of its hexameric ring structures, and into the proteolytic barrel of ClpP (33, 34). Thus, by homology, it has been hypothesized that ClpB and Hsp104 must employ the same mechanisms as ClpA. However, this has been difficult to test, in part, because of the lack of covalent modification coupled to ClpB/Hsp104 catalyzed processing of protein substrates. This means that when ClpB/Hsp104 processes a protein substrate, they do not hand it off to an associated protease. Consequently, the substrate both enters and leaves the reaction without covalent modification.

### Evidence of processive protein unfolding and translocation catalyzed by ClpB

In acknowledgment of the difficulty in detecting unfolding and translocation without covalent modification, Weibezahn et al embarked on a clever protein engineering strategy to test for threading through the axial channel of ClpB (35). In their strategy they constructed a variant of ClpB that included the IGL loop from ClpA. The IGL loop is responsible for the interaction between ClpA and ClpP. The rationale being that if ClpB was “forced” to interact with the protease, ClpP, and if proteolytic fragments were detected then this must indicate that ClpB threads substrate through the axial channel and into ClpP for proteolysis in the same way as ClpA.

Weibezahn et. al. reported mixing the intrinsically disordered protein, α-casein, ClpB with the IGL loops added (ClpB(IGL)), ClpP, and ATP and observed the loss of the band representing α-casein on an SDS-PAGE gel (35). However, using their construct, we showed faster loss of full length α-casein when this experiment was performed in the absence of ATP or any other energy source (17), a control that was missing in the Weibezahn et al paper.

Cleavage in the absence of ATP calls into question what is being detected with the ClpB(IGL) construct since one expects enzyme catalyzed protein translocation and subsequent threading into ClpP to be an energy requiring process. Indeed, degradation is observed in the presence of ATP but it is observed to be faster in the absence of ATP. Thus, one is left asking what proportion of the observed degradation is energy dependent vs. energy independent. Equally important, the loss of the α-casein band on a gel can occur from a single cleavage event. Thus, loss of the band does not reveal if ClpB processively threaded the substrate into ClpP and multiple cleavage events occurred or if only one cleavage event occurred.

Wickner and co-workers have shown that both ClpB and Hsp104 can induce a loss of GFP fluorescence when GFP contains the first seventy amino acids of the RepA protein at the N-terminus as a binding site (14). With those results in mind, we attempted to develop the stopped-flow approach reported here using RepA-GFP. However, we observed loss of fluorescence upon simply mixing ClpB/Hsp104 with RepA-GFP and ATPγS (data not shown). Under the impression that ATPγS would only support assembly and binding but not ClpB catalyzed protein unfolding, we interpreted those observations to indicate that, in the presence of only ATPγS, ClpB bound to the junction and melted sufficient secondary structure to result in cooperative unfolding of GFP and subsequent loss of fluorescence. After all, protein unfolding is often cooperative, thus the motor may only need to destabilize some of the folded region of GFP to induce complete unfolding (36). This led to questions about how we would determine the extent to which the structured region must be unfolded before GFP fluorescence is extinguished in a single turnover experiment. Thus, the RepA-GFP construct was abandoned.

### Development of transient state kinetics approach to examine the elementary steps in enzyme catalyzed protein unfolding and translocation

Our objective has been to develop a single turnover stopped-flow method that would report on the elementary kinetic steps in a single-round of enzyme catalyzed protein unfolding and translocation. We have been seeking to quantify the elementary rate constants, kinetic step-sizes, and processivity for the protein disaggregating machines, ClpB and Hsp104. To this end, we previously applied the single turnover stopped-flow approach that used unfolded polypeptides ranging in length between 30 and 50 amino acids as well as truncations of the unstructured protein α-casein of lengths up to 127 amino acids. Based on the observations of Doyle et al (14) we used a mix of ATP and ATPγS to activate the motor as we have done here (17, 18, 25).

Importantly, we define protein unfolding and translocation as two kinetically distinct processes. Thus, when using unstructured polypeptide chains we interpret the results to represent polypeptide translocation and not enzyme catalyzed protein unfolding. Using the stopped-flow approach with unstructured polypeptides we observed that both ClpB and Hsp104 proceeded through two rate-limiting steps before rapid dissociation, independent of the substrate length provided (17, 18). From those observations we proposed that ClpB and Hsp104 are non-processive translocases on unstructured polypeptides, P ∼ 0.6, see Equation 2. However, we acknowledged, for both enzymes that we could not rule out the possibility that we were only detecting two slow steps either before or after fast translocation that was outside the temporal resolution of the stopped-flow, i.e. millisecond resolution.

Avellaneda *et. al*. reported fast and processive translocation of mechanically unfolded Maltose Binding Protein (MBP) using optical tweezer measurements and a “hyper-active” variant of ClpB that does not require ATPγS to be activated (13, 19). In their approach they attached the N-and C-terminus of MBP to polystyrene beads and optically trapped each bead. The protein was then mechanically unfolded by increasing the distance between the two optically trapped beads and then 1 μM ClpB(Y503D) was added. Upon adding ClpB(Y503D) and ATP they report rapid reduction in the distance between the two beads. From this measurement, they report ClpB translocates on the unfolded polypeptide chain at rates of ∼250 and 450 aa s^-1^. As we have expressed elsewhere, it is not clear if these rates represent the activity of a single ClpB hexamer (20). However, if correct, the values should be interpreted as a rate of translocation on an unfolded polypeptide chain and not a rate for protein unfolding.

Here we report a rate of ∼3.1 and 0.9 aa s^-1^ for enzyme catalyzed protein unfolding of tandem repeats of the Titin I27 domain in the presence of 500 μM ATP and either 150 or 500 μM ATPγS, respectively. These rates are two orders of magnitude slower than what was reported by Avellaneda *et. al*. (19), consistent with translocation and protein unfolding being kinetically distinct processes. Indeed, the measurement here are in the presence of ATPγS but the increase in rate from ∼0.9 to 3.1 with a reduction in ATPγS by a factor of 3.3 is not suggesting that the rate will increase by two orders of magnitude if ATPγS was absent. Rather, we hypothesize that what we are detecting is rate-limited protein unfolding followed by undetectably fast translocation up to the next folded region and then the process repeats. However, whether or not the rates of translocation between unfolding events are as fast as reported by Avellaneda et al requires further testing (20).

The rate-limiting protein unfolding of the Titin I27 domains followed by rapid translocation has not been reported for ClpAP and ClpXP. Olivares et al, using optical tweezer approaches with similar Titin I27 domains used here, reported rate limiting translocation after fast protein unfolding for both ATP dependent proteases, ClpAP and ClpXP (37). They observed a dwell time preceding protein unfolding of ∼0.9 and ∼0.8 s for ClpXP and ClpAP, respectively. The inverse of this can be taken as the rate constant for protein unfolding and would yield a value of ∼1.2 s^-1^, which is in good agreement with our observed rate constant of ∼0.9 – 3 s^-1^ depending on the ATP:ATPγS mixing ratio. For ClpB, we propose that the slow unfolding is followed by rapid translocation on the unfolded chain where translocation by ClpB must be much faster than for ClpAP and ClpXP.

Importantly, using single turnover stopped-flow experiments we showed that ClpAP and ClpA exhibited different translocation mechanisms including overall rate, rate constants, and kinetics step-sizes on unfolded polypeptide chains. We interpreted those differences to indicate that ClpP allosterically impacts the translocation mechanism employed by ClpA (38, 39). Thus, ClpA and ClpAP should be considered different enzymes. Consequently, it is not surprising that ClpA and ClpB would exhibit marked differences in their mechanisms of protein unfolding and translocation.

### Interpretation of the kinetic step-size

From the model independent peak time analysis, we determined an overall rate of protein unfolding. However, the rate of protein unfolding is a convolution of the number of amino acids unfolded per step and the rate constant defining that step. From global analysis we can deconvolute the rate constant and step-size from the overall rate thereby extracting additional information about the elementary mechanism.

From global fitting of the stopped-flow time courses to an *n*-step sequential mechanism we report a kinetic step-size of ∼57 and ∼58 amino acids per step at 1:1 and ∼3:1 ATP:ATPγS, respectively. Importantly, the kinetic step-size represents the average number of amino acids unfolded or translocated between two rate-limiting steps. In contrast to the kinetic step-size, we would define a mechanical step-size as the physical distance the motor translocates on the polypeptide chain.

In some cases, the kinetic step-size and mechanical step-size may be the same. For example, we reported a kinetic step size for ClpAP to be ∼5 amino acids per step (38). From optical tweezer experiments it was later reported that ClpAP exhibited a mechanical step-size of 4-8 amino acids per step (40), indicating that the same process was being monitored in both experimental approaches.

Here we conclude that it is unlikely that ClpB physically traverses 56 – 58 amino acids in a single step. So, what is the meaning of this kinetic step-size? The magnitude must be governed by the structural properties of both the motor and the substrate being unfolded and translocated. From cryo-EM structures of ClpB bound to unfolded polypeptide chains, ∼ 26 amino acids span the axial channel of the hexameric ring (41). It seems unlikely that the motor would traverse 26 amino acids in a single step but the full length of the axial channel or 26 aa could be considered an upper limit on mechanical stepping. Nevertheless, the large kinetic step-size observed here, taken with a substantially slower rate on a folded compared to an unfolded polypeptide, suggests rate-limited protein unfolding of ∼60 amino acids followed by rapid translocation that is outside of the millisecond temporal resolution of the stopped-flow. That is to say ClpB must apply force to the folded structure that results in the cooperative collapse of ∼60 amino acids or nearly a full Titin I27 domain of 98 amino acids followed by rapid translocation up to the next folded domain, see Fig. 5.

A mechanical translocation step-size of 2 amino acids per step has been proposed from many cryo-EM structures of AAA+ motors bound to polypeptide substrates (42, 43). This is based on the distance between polypeptide contacts across the axial channel of the motor. However, to our knowledge, an experimental determination of a mechanical step-size as small as two amino acids per step has yet to be reported. Nevertheless, rate-limiting protein unfolding of ∼60 amino acids followed by fast translocation in small sub-steps remains a model to be tested.

### Quantification of unfolding vs. translocation processivity

The conclusion that ClpB is a non-processive translocase on unfolded polypeptide chains was based on the lack of any detectible length-dependence in the time-courses (17). The collected time courses were most consistent with two-step dissociation. Here, we have shown that ClpB is able to unfold up to three tandem repeats of stably folded Titin I27 domains with a total substrate length of 362 amino acids. This indicates that, after unfolding the substrate, ClpB exhibits a translocation processivity of at least N = 362 amino acids, where N is defined as the processivity in the number of amino acids translocated per binding event.

The processivity, N, in number of amino acids is related to the processivity as a probability, P, as defined in Eq. 3, where m is the step-size (32).

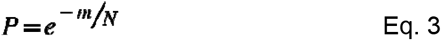

Since the relationship between the probability processivity, P, and the number processivity, N, requires knowledge of the step-size, m, we can approximate the processivity, P, for translocation in the presence of fold to be in the range of P = 0.994 to P = 0.946 for a step-size between 2 and 20 amino acids, respectively, since we do not know the translocation step-size with certainty. In contrast, if we use a step-size of ∼60 amino acids, which we interpret as a protein unfolding step-size, in Eq. 3 then P = ∼ 0.85. Thus, these differences in processivity may suggest that when ClpB is challenged by stably folded regions the processivity for this process is reduced. Thus, going forward, it will be interesting to determine how processivity relates to stability of the folded regions.

One explanation for the apparent disparity between this study and our previous work is that the previous lack of structure in the substrate may serve to signal ClpB to dissociate. If this were the case then after ClpB unfolds the last ∼60 amino acids and no more folded structure is down stream of the motor then one would expect ClpB to dissociate. However, given that the PIFE distance is 0 – 3 nm (44), detecting PIFE is consistent with ClpB being within ∼12 amino acids of the C-terminus before dissociation. This predicts that some number of translocation steps do occur on the unstructured polypeptide after complete unfolding of the final Titin I27 domain.

Another explanation is because ATPγS supports processive protein unfolding, it is possible that ATPγS driven translocation obscured the observation of a length-dependence in our previous studies. If ClpB was coupling ATPγS hydrolysis to processive rounds of translocation during the pre-incubation time on the stopped-flow then one would expect that, upon rapid mixing with ATP, the motor proteins would be randomly distributed on the polypeptide substrates. We know from studies on ssDNA translocases that random binding to the lattice to be translocated will still lead to the observation of a length dependence in a single turnover stopped-flow experiment (45, 46). Thus, if randomly bound motor on the peptides was the state of the system upon rapid mixing with ATP then we would predict the observation of a length dependence that we did not previously detect (17).

Alternatively, in the pre-incubation syringe, if ClpB used ATPγS to translocate up to the fluorophore and stalled at the end until rapid mixing with ATP that induced dissociation, then this situation could lead to the apparent non-processive translocation behavior previously reported. If this model is correct then the dissociation rate constant detected here, *k*_*end*_, should be equivalent to the reported rate constants for dissociation from unstructured substrates. Here we observe dissociation from the end to range from *k*_*end*_ = 0.01 – 0.15 s^-1^ at 1:1 ATP:ATPγS and from our previous report we detected dissociation from the unstructured substrates to be between 0.03 – 0.15 s^-1^. These numbers are in good agreement and may indicate that, in our previous study, we only detected dissociation from the end due to a stalled motor at either the N-or C-terminus. However, the experiments performed in the presence of only ATPγS, shown in Fig. 2 B, show a loss in signal consistent with the eventual dissociation of ClpB after arrival at the fluorophore. Thus, the time-courses reported here are consistent with slow dissociation from the end.

We propose that the best explanation is that ClpB rapidly translocates on unfolded polypeptide chains so fast that, on unfolded substrates, we only detect two slow dissociation steps. With the agreement between the rates of dissociation from the end of the substrates reported here and the values previously reported from unstructured polypeptides,(17) we favor a model where rapid translocation is followed by slow dissociation from the end, see k_end_ in Fig. 5. This interpretation is most consistent with recent reports on unfolded polypeptides (19, 21, 22).

We interpret the results reported here to indicate that protein unfolding is rate limiting, see k_u_ in Fig. 5, and ∼60 amino acids are unfolded between two rate limiting steps at the ATP/ATPγS concentrations used. Each Titin I27 domain is 98 amino acids so a step-size of ∼60 amino acids is less than one full domain. But a complete examination of the ATP concentration dependence of the kinetic step-size is underway. Thus, it remains possible that at elevated ATP more structure could be unfolded. Nevertheless, it is interesting to note that Oberhauser et al reported that tandem repeats of the Titin I27 domain were unfolded in steps of 22 nm when pulled using AFM experiments (47). If one uses 0.34 nm per amino acid then the AFM experiments reveal ∼65 amino acids unfolded per step in good agreement with the kinetic step-size reported here.

More substantial work is needed to fully understand the mechanisms of ClpB catalyzed protein unfolding and translocation. Several methods have been developed to examine the reactions catalyzed by ClpB in the absence of the co-chaperones, KJE. Here we have used a mixture of ATP and ATPγS, which is one of the methods (14). However, applying our approach to the hyper-active variants is of interest, e.g. ClpB(Y503D). In addition, further development of the sequential mixing approach that will allow us to include the full complement of co-chaperones, KJE, is underway (25).

Here we interpret our results to indicate that protein unfolding is rate-limiting and translocation is fast, see Fig. 5. This predicts differences in the rate and kinetic step-size depending on the stability of the fold presented to the motor. With the establishment of the sequential mixing approach developed here we are well positioned to address this question.

Moreover, we are now positioned to address questions on directionality and processivity that have been out of our reach for motors like ClpB and Hsp104. Further, by interrogating the impact of protein stability on mechanical unfolding rates, rate constants, and step-sizes we may acquire insights into protein folding that complements common techniques such as chemical denaturation and heat denaturation. Moreover, how ATP hydrolysis is coupled to the reaction detected and the role of Domain 1 and Domain 2 ATP binding and hydrolysis sites remain to be interrogated. Nevertheless, the work presented here has opened the door to answering these questions and many more about the fundamental reactions catalyzed by protein disaggregating machines and their role in protein homeostasis. With that in mind, so long as one can define a binding sequence, this approach can be broadly applied to the many AAA+ molecular motors that do not modify the substrates they process.

## Materials and Methods

### Buffers and reagents

Buffers were prepared with reagent-grade chemicals using 18 MΩ deionized water from a Purelab Ultra Genetic system (Evoqua, Warrendale, PA). Buffer H200 contains 25 mM HEPES, pH = 7.5 at 25 °C, 10 mM MgCl_2_, 200 mM NaCl, 2 mM 2-mercaptoethanol and 10 % (v/v) Glycerol. *E. coli* ClpB was purified as described (30). All ClpB concentrations are reported in monomer units. ATP and ATPγS were purchased from Thermo Fischer Scientific (Waltham, MA) and CalBiochem (La Jolla, CA), respectively. Both ATP and ATPγS were dialyzed into H200 using 100-500 Da molecular weight cut-off dialysis tubing (Thermo Fischer Scientific, Waltham, MA). α-casein was purchased from Sigma-Aldrich (Darmstadt, Germany), dissolved in 6 M Guanidine hydrochloride, 20 mM HEPES, pH 7 at 25 °C, and dialyzed into H200 using 10 kDa molecular weight cut-off dialysis tubing (Thermo Fischer Scientific, Waltham, MA).

### Purification of RepA-Titin_x_

The His_6_-RepA-Titin_X_ proteins are composed of an N-terminal 6 His tag with a thrombin cleavage sequence for tag removal (MGSSHHHHHH SSGLVPRGSH). The 6 His tag is followed by the first 70 amino acids of the phage P1 RepA protein (MNQSFISDIL YADIESKAKE LTVNSNNTVQ PVALMRLGVF VPKPSKSKGE SKEIDATKAF SQLEIAKAEG) (24). For RepA-Titin_1_ the RepA sequence is directly connected to the Titin I27 domain (PDB ID: 2RQ8)(25) with all native cysteine residues changed to alanine and the final sequence is given by MLIEVEKPLY GVEVFVGETA HFEIELSEPD VHGQWKLKGQ PLAASPDAEI IEDGKKHILI LHNAQLGMTG EVSFQAANT KSAANLKVKE L. Additional Titin I27 domains are connected with an eight residue linker, RSKLGTRM. Each construct contains a single cysteine at the C-terminus for fluorescent modification. The genes encoding for RepA(1-70)Titin_X_ were constructed and cloned into the pET28a vector by Genscript (Piscataway, NJ). The total substrate lengths are 168, 265, and 362 amino acids for RepA-Titin_1_, RepA-Titin_2_, RepA-Titin_3_, respectively.

The pET28a plasmids were transformed into ΔClpP_WT_-BL21 cells using electroporation (48). ClpP_WT_ knockout BL21(DE3) cells, ΔClpP_WT_-BL21, were constructed using recombineering (49). ClpP_WT_ gene in BL21(DE3) genome was substituted with an Ampicillin cassette similar to what has been done earlier with ΔClpA_WT_-BL21(DE3) cells (50). The ΔClpP_WT_-BL21 cells were grown in LB Miller growth media (Thermo Fischer Scientific, Waltham, MA) containing 100 μg/mL Ampicillin.

ΔClpP_WT_-BL21 cells were grown to mid-log growth and induced with 1 mM IPTG to express His_6_-RepA-Titin_x_. The cell paste was suspended in cell lysis buffer [40 mM Tris pH 7.5 at 4 ºC, 20 mM Imidazole, 500 mM NaCl, 2 mM 2-mercaptoethanol, 10 % (w/v) sucrose, 20 % (v/v) glycerol]. Cells were lysed using a French Press at 4 ºC. DNase I (Thermo Fischer Scientific, Waltham, MA) and RNase A (Thermo Fischer Scientific, Waltham, MA) were added to the lysate and incubated at 4 ºC for 15 minutes. Cell debris was pelleted by centrifugation for 120 minutes at 28,000 g and 4 °C in a ThermoScientific Fiberlite F14-6×250 rotor. His_6_-RepA-Titin_x_ was in the soluble fraction of the lysate.

Isolation of His_6_-RepA-Titin_x_ started with batch purification. The soluble fraction was added to HisPur™ Ni-NTA resin (Thermo Fischer Scientific, Waltham, MA) and incubated with rocking for 75 minutes at 4 ºC. The resin was subsequently washed with 5-10 column volumes of binding buffer (40 mM Tris, pH 7.5 at 4 ºC, 20 mM Imidazole, 500 mM NaCl, 2 mM 2-mercaptoethanol, 10 % (v/v) glycerol). The resin was then incubated with 1 column volume of elution buffer (40 mM Tris, pH 7.5 at 4º C, 500 mM Imidazole, 500 mM NaCl, 2 mM 2-mercaptoethanol, 10%(v/v) glycerol) for 1 hour at 4º C on a rocker. The slurry, comprising the resin and the elution buffer was altogether poured into a gravity column and incubated for 10 minutes to form a resin bed. His_6_-RepA-Titin_x_ bound to the resin was expected to elute with the elution buffer i.e. under high Imidazole conditions. Fractions of 1 mL were collected from the gravity column. Further the resin bed was washed with 5-10 column volumes of elution buffer to elute any remaining His_6_-RepA-Titin_X_ from the resin. Fractions containing His_6_-RepA-Titin_X_ were identified using a NuPAGE™ gel (Thermo Fischer Scientific, Waltham, MA) with Coomassie staining. The fractions of interest were dialyzed into Thrombin digestion buffer (20 mM Tris, pH 7.5 at 4 ºC, 150 mM NaCl, 2.5 mM CaCl_2_) using a 10 kDa cutoff dialysis tubing (Thermo Fischer Scientific, Waltham, MA).

His_6-_RepA-Titin_x_ was subjected to Thrombin protease (Sigma Aldrich, Burlington, MA) following the manufacturer’s protocol. The protein sample was passed through His Trap™ FF crude column (GE Healthcare, Piscataway, NJ). We expected the undigested protein to remain bound to the column. After washing the column with the digestion buffer, RepA-Titin_x_ was washed out with the binding buffer. All the fractions of interest were pooled together and dialyzed into a low salt buffer (20 mM Tris, pH 7.5 at 4 ºC, 20 mM NaCl, 2 mM 2-mercaptoethanol). Further, the protein sample was passed through a Hi-Trap Q-Sepharose FF crude (GE Healthcare, Piscataway, NJ) column. The column was washed with a linear gradient of low salt and high salt buffer (20 mM Tris, pH 7.5 at 4 ºC, 1 M NaCl, 2 mM 2-mercaptoethanol) starting with 100% low salt buffer (20 mM NaCl) and moving gradually to 100% high salt buffer (1 M NaCl). RepA-Titin_x_ eluted at low salt conditions. The fractions of interest were dialyzed into labeling buffer (20 mM HEPES, pH= 7.5 at 4 ºC, 50 μM TCEP).

Labeling of C-terminal Cysteine on RepA-Titin_x_ was carried out using a C_2_-maleimide reaction with Alexa Fluor™ 555 fluorophore (Invitrogen, Waltham, MA) following the manufacturer’s protocol except 3 (fluorophore) : 1 (protein) molar excess. After labeling, RepA-Titin_x_-Alexa Fluor 555 along with free Alexa Fluor 555 was dialyzed into the H200 buffer. Free Alexa Fluor 555 was then separated from RepA-Titin_x_-Alexa Fluor 555 using a HiPrep 26/10 desalting column (GE Healthcare, Piscataway, NJ). The labeled protein sample was dialyzed into storage buffer (40 mM Tris, 500 mM NaCl, 2 mM 2-mercaptoethanol, 10%(v/v) glycerol, 2 mM EDTA) and stored at -80 ºC. The labeling efficiency of RepA-Titin_x_-Alexa Fluor 555 in H200 buffer was observed to be 65-100%. The reported concentrations of RepA-Titin_x_ substrates are determined spectrophotometrically by measuring the absorbance of Alexa Fluor 555 at 555 nm and using an extinction coefficient of 158 000 M^-1^ cm^-1^.

### Structures of RepA-Titin_x_

The raw structures of RepA-Titin_X_ with X = 1, 2, or 3 obtained from simulations using AlphaFold are shown in the supplemental materials (51). Starting from the N-terminus, the first 70 amino acids represent the N-terminus of the Phage P1 RepA protein. AlphaFold predicted low confidence on α helices present in the RepA 1 – 70 sequence, see supplemental. This is consistent with previous reports that this sequence is not likely structured (23, 52, 53). Thus, we used ‘Sculpting’ function in Schrodinger (54, 55) to unfold the α helices in the RepA sequence in the AlphaFold predicted structures to yield the structures shown in Fig. 1 A. The TitinI27 regions are predicted to have folded β sandwich structure consistent with the published structure (PDB ID: 2RQ8) (56). The connectors between tandem repeats of TitinI27 are predicted to be unstructured as shown in both supplemental materials and Fig. 1. Cysteine is shown in space-filling at the C terminus of each substrate. Alexa Fluor 555 is attached to the C-terminal Cysteine, not shown.

### Standard mixing stopped-flow experiments

ClpB and RepA-Titin_X_ were dialyzed into H200 buffer using 50 kDa and 10 kDa molecular weight cutoff dialysis tubing, respectively. The experiments were performed on an SX20 Applied Photophysics stopped-flow fluorometer (Leatherhead, U.K.) under standard mixing set-up at 25 °C as shown in Fig. 1 C.

1.5 μM ClpB was incubated for 5 minutes with 300 μM ATPγS to form the active hexameric complex and subsequently incubated for 10 minutes with 100 nM RepA-Titin_x_ substrate to form the pre-bound complex. The pre-bound complex was loaded into Syringe 1 and 400 μM ATP and 20 μM α-casein into Syringe 2 of the apparatus. All the components are diluted two-fold upon mixing. Before collection of the first shot, the observation channel was thoroughly washed with the solutions of each syringe to equilibrate the instrument. For comparing the relative increase in fluorescence signal across RepA-Titin_x_ substrates, processing of raw time-courses were done using Eq 4.

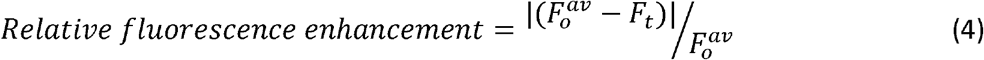

where 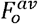 is the average of the first few constant raw fluorescence data points and F_t_ is raw fluorescence signal at a given time.

### Sequential mixing stopped-flow experiments

8 μM ClpB was rapidly mixed with 2000 μM ATPγS and 200 nM RepA-Titin_x_ substrate. After mixing, the reagents incubate in the ageing loop for a user-defined period of time, Δt_1_. 1000 μM ATP and 40 μM α-casein in Syringe 3 were rapidly mixed with the pre-bound complex of ClpB with RepA-Titin_x_ formed during sequential mixing in ageing loop, see Fig. 3 A. All the components in Syringes 1 and 2 are diluted four-fold when present in the observation chamber while components in Syringe 3 are diluted two-fold. In the observation channel, Alexa Fluor 555 is excited at 555 nm and the emission signal is collected using a 570 nm long pass filter.

### Determination of reduced length

For each set of time-courses collected using RepA-Titin_X_, we plot total length versus peak time. The intercept of this plot is C, which is the sum of pre-translocated distance with ATPγS and excluded length. Excluded length is defined as the sum of dangle distance and occluded length, see Supp. Fig. 5. So, we subtract *C* from total length, *L* to get reduced length of substrate, *L’. L’* is the available length for ClpB to translocate and unfold. To note, *C* is determined individually for each replicate at a particular Δt_1_. The average values of *C* at different Δt_1_’s are shown in Fig. 3 E and Supp. Fig. 7 D.

### Model dependent analysis of the experimental time-courses

The time-courses were fit using the custom built MATLAB(Mathworks, Natick, MA) toolbox, MENOTR(57). Eq 5 described the data set for each RepA-Titin_x_ and the resultant kinetic parameters are described in table 2. Out of all parameters, *k*_*U*_ and *m* were fit globally across each RepA-Titin_X_.

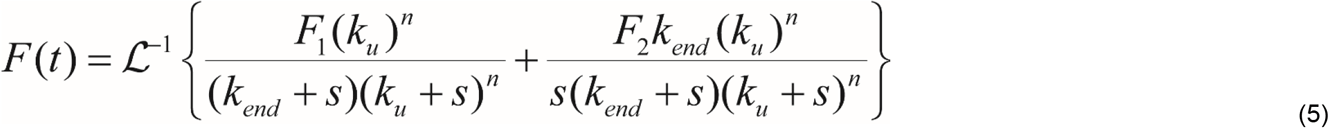

Where *F*_*1*_ and *F*_*2*_ are two fluorescence amplitudes for the last intermediate, I_(L-nm)_ and unfolded RepA-Titin_x_ respectively, *s* is the Laplace variable and other kinetic parameters are defined as per scheme 1, see Fig. 4 A.

## Acknowledgments

We thank Elizabeth Duran, Clarissa Durie, and members of the Lucius lab for their critical discussions of the results and the manuscript. This work was supported by the National Science Foundation (grant NSF MCB-1412624 to A.L.L.). Computational work was performed using the UAB High Performance Computing (HPC) Cheaha, which is supported in part by the National Science Foundation under Grants No. OAC-1541310, the University of Alabama at Birmingham, and the Alabama Innovation Fund.

## Figures and Tables

**Scheme 1.**
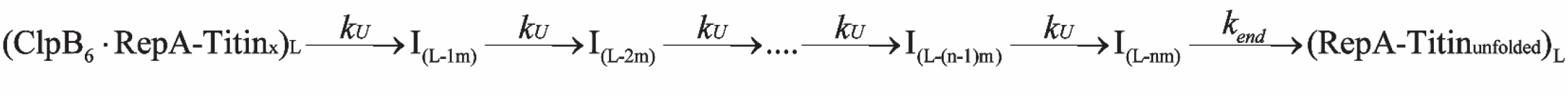

