## Supplemental Figures for "Single Turnover Transient State Kinetics Reveals Processive Protein Unfolding Catalyzed by *Escherichia coli* ClpB"

### Supplemental Figure 1.

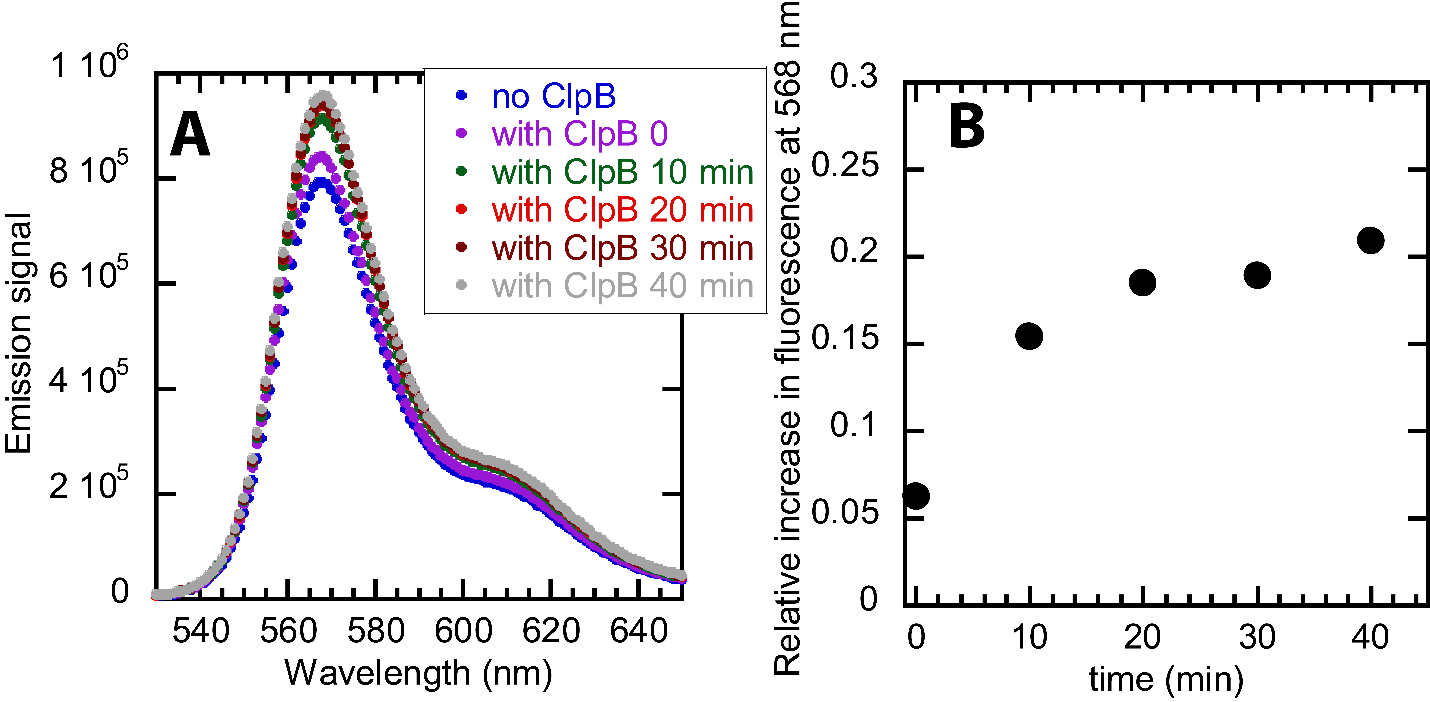

Supplemental Figure 1

Testing Protein Induced Fluorescence Enhancement with an unstructured substrate of 50 amino acids

A) 2 μM ClpB in the presence of 1 mM ATPγS was incubated with 0.5 μM polypeptide labeled with AF555. The emission spectra by exciting AF555 at 520 nm were collected every 10 minutes. The blue trace represents emission spectra collected with no ClpB. The purple trace represents emission spectra collected immediately after mixing ClpB with ATPγS and polypeptide. 10 min(green), 20 min(red), 30 min(brown), and 40 min(grey) represent the incubation times of ClpB with ATPγS and polypeptide.
B) Relative fluorescence enhancement was calculated at 568 nm by using Eq 4 and is plotted as a function of incubation times of ClpB with ATPγS and polypeptide. With the increase in incubation times, we observe fluorescence enhancement upon binding of ClpB with polypeptide substrate.

### Supplemental Figure 2.

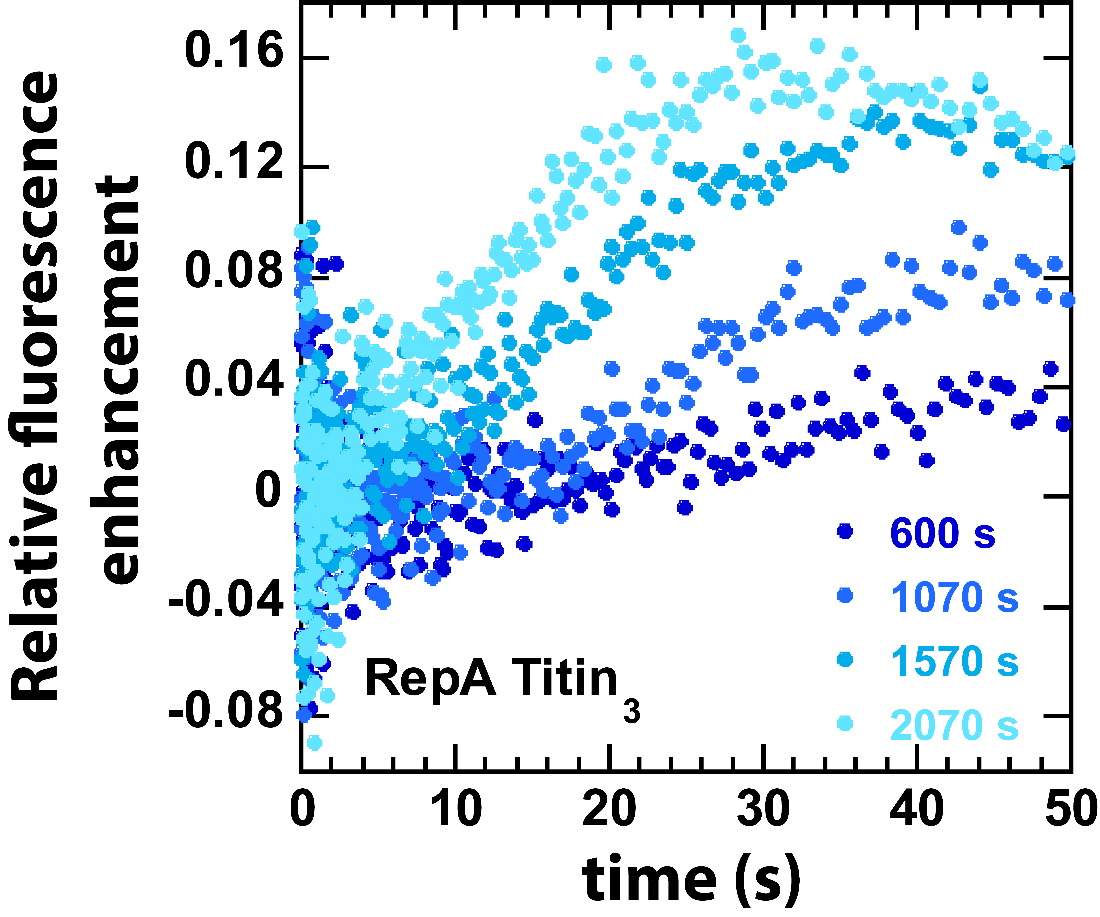

Supplemental Figure 2: A zoomed-in version of Fig. 1 E to show the decreasing lag with increasing incubation time of the pre-bound complex.

### Supplemental Figure 3.

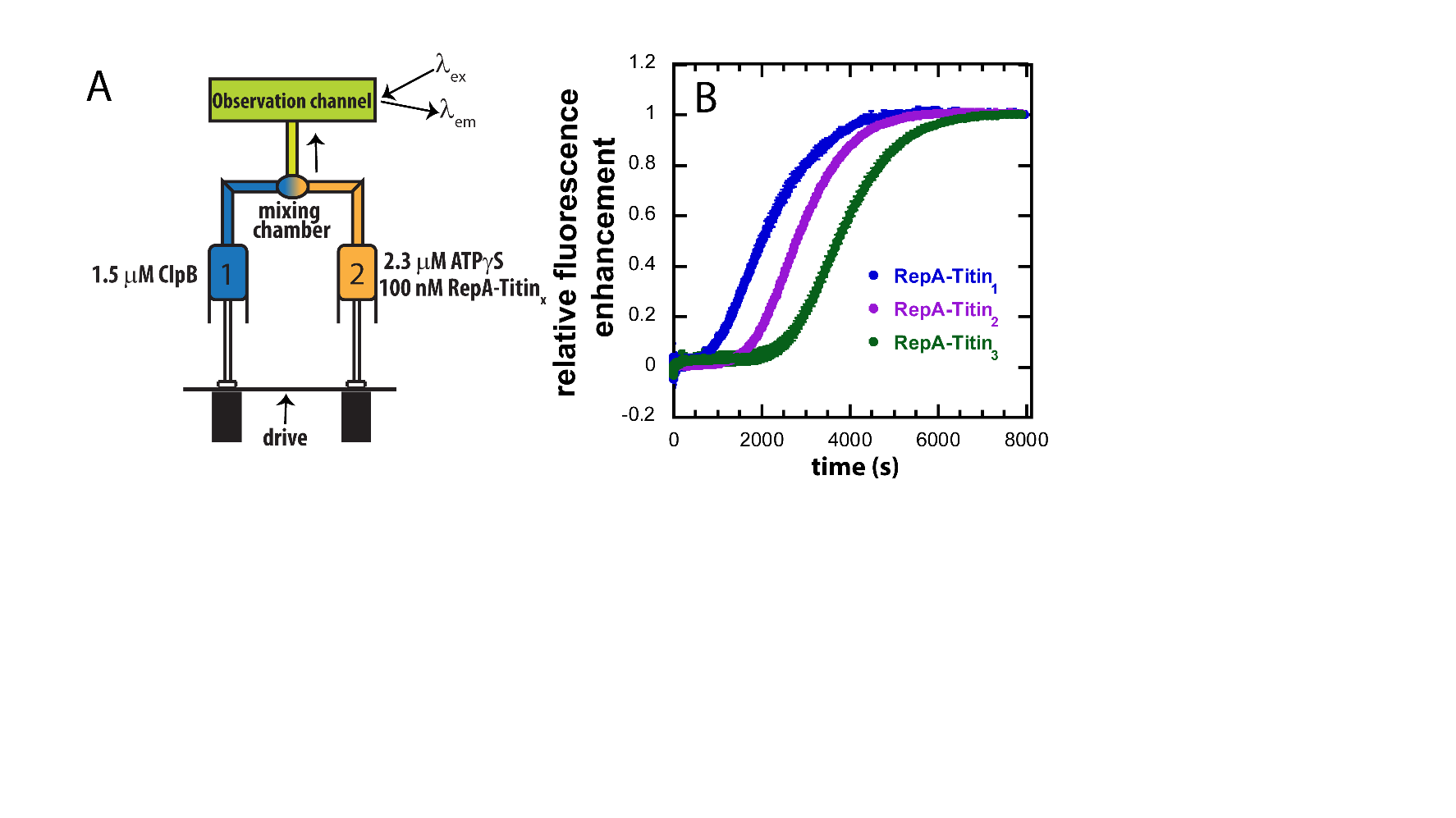

Supplemental Figure 3: 1.5 μM ClpB(Syringe 1) is mixed with 2.3 μM ATPγS (Syringe 2) and 100 nM RepA-Titin_X_ (Syringe 2), where x =1, 2, 3, using standard mix on stopped-flow apparatus. Under these conditions, upon mixing, ClpB binds ATPγS, assembles into hexamers competent for binding RepA- Titin_1_ and proceeds to unfolding. In this setup, the fluorescence change is monitored over time. The scatter plot shows the experimental time-courses with the maximum fluorescence normalized to one obtained using three different RepA-Titin_X_ substrates. The experimental time-courses are an average of at least 3 or more successive time-courses and the standard deviation of these averaged time-courses are shown as error bars on the data points.

### Supplemental Figure 4.

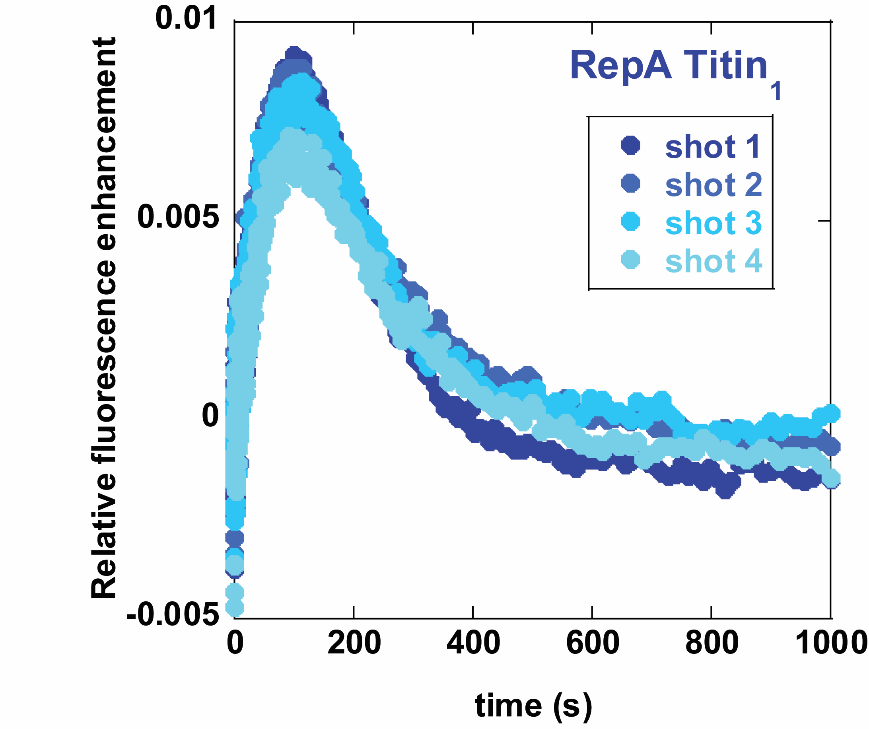

Supplemental Figure 4: Reproducibility using sequential mixing strategy. Successive time-courses for RepA-Titin_1_ were collected using the schematic in Fig. 3 A. Shots 1 – 4 indicates time-courses collected in succession.

### Supplemental Figure 5

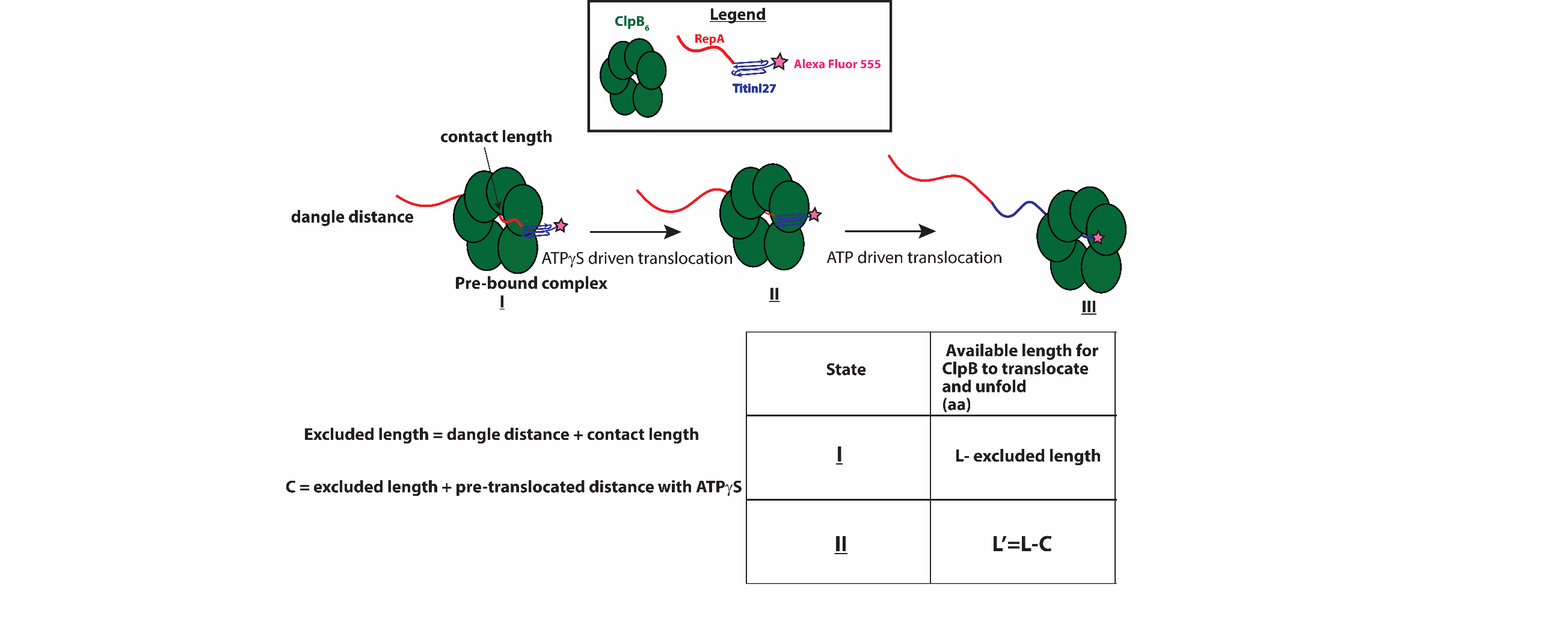

Supplemental Figure 5:

Cartoon representation showing translocation and unfolding by ClpB on RepA-Titin_1_ from states I through III: (I) Pre-bound complex of ClpB (green) with RepA-Titin_1_ has dangle distance and contact length unavailable for translocation and unfolding. (II) After ATPγS driven translocation has occurred, the unavailable length is the pre-translocated distance and excluded length. (III) On ATP driven translocation, ClpB completely unfolds and translocates on RepA-Titin_1_.

### Supplemental Figure 6.

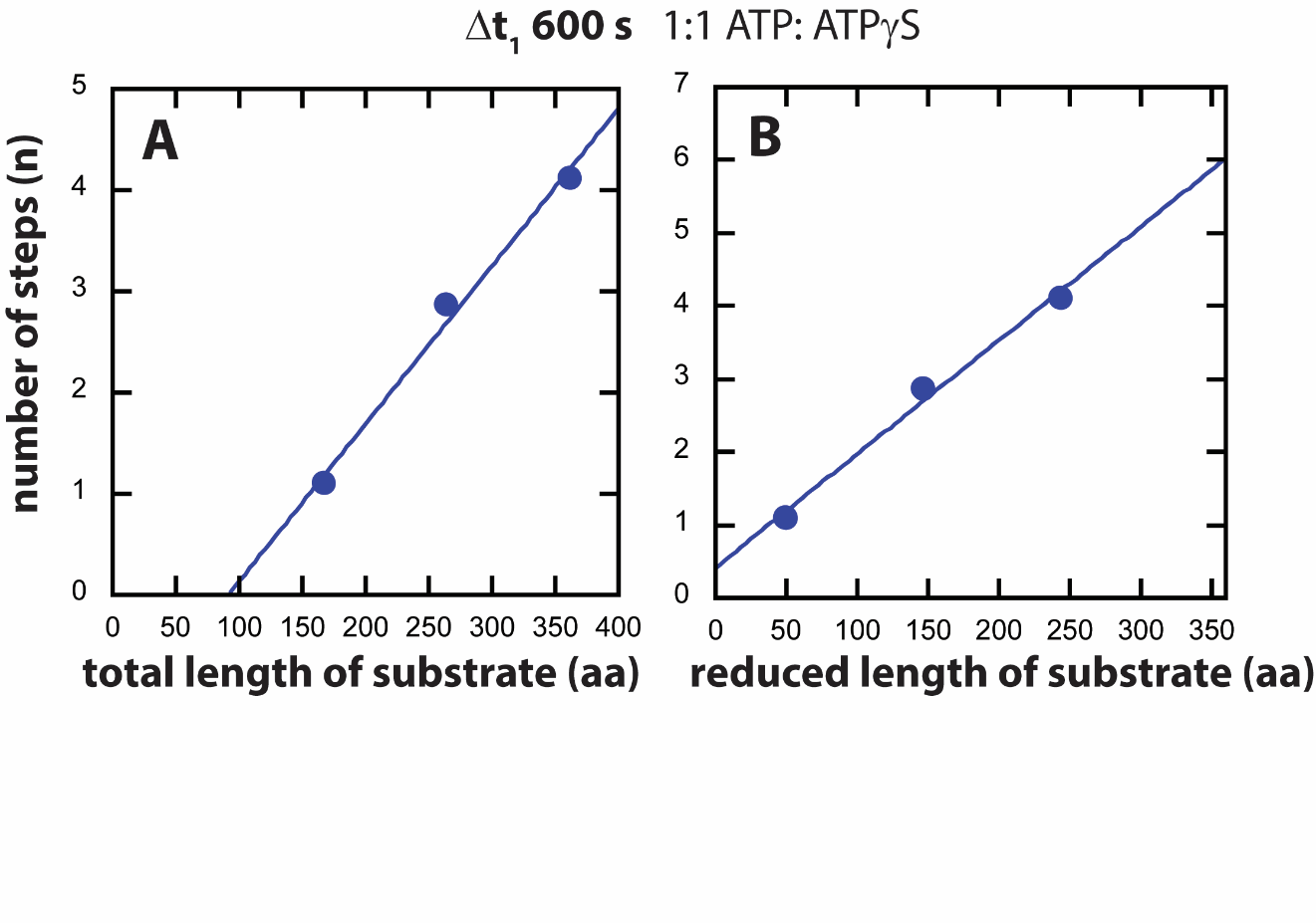

Supplemental Figure 6: Determination of number of steps, *n*, from fitting the time-courses represented in Fig. 3 B to Scheme 1. The number of steps, *n*, (shown in circles) is plotted as a function of (A) number of steps vs total length of substrate. Solid line yields a slope = (0.016 ± 0.002) aa^-1^ and intercept = (-1.4 ± 0.4) steps. (B) Reduced length of substrate calculated by subtracting *C* = 118 aa from the length. Solid line yields a slope = (0.016 ± 0.002) aa^-1^ and zero intercept.

### Supplemental Figure 7.

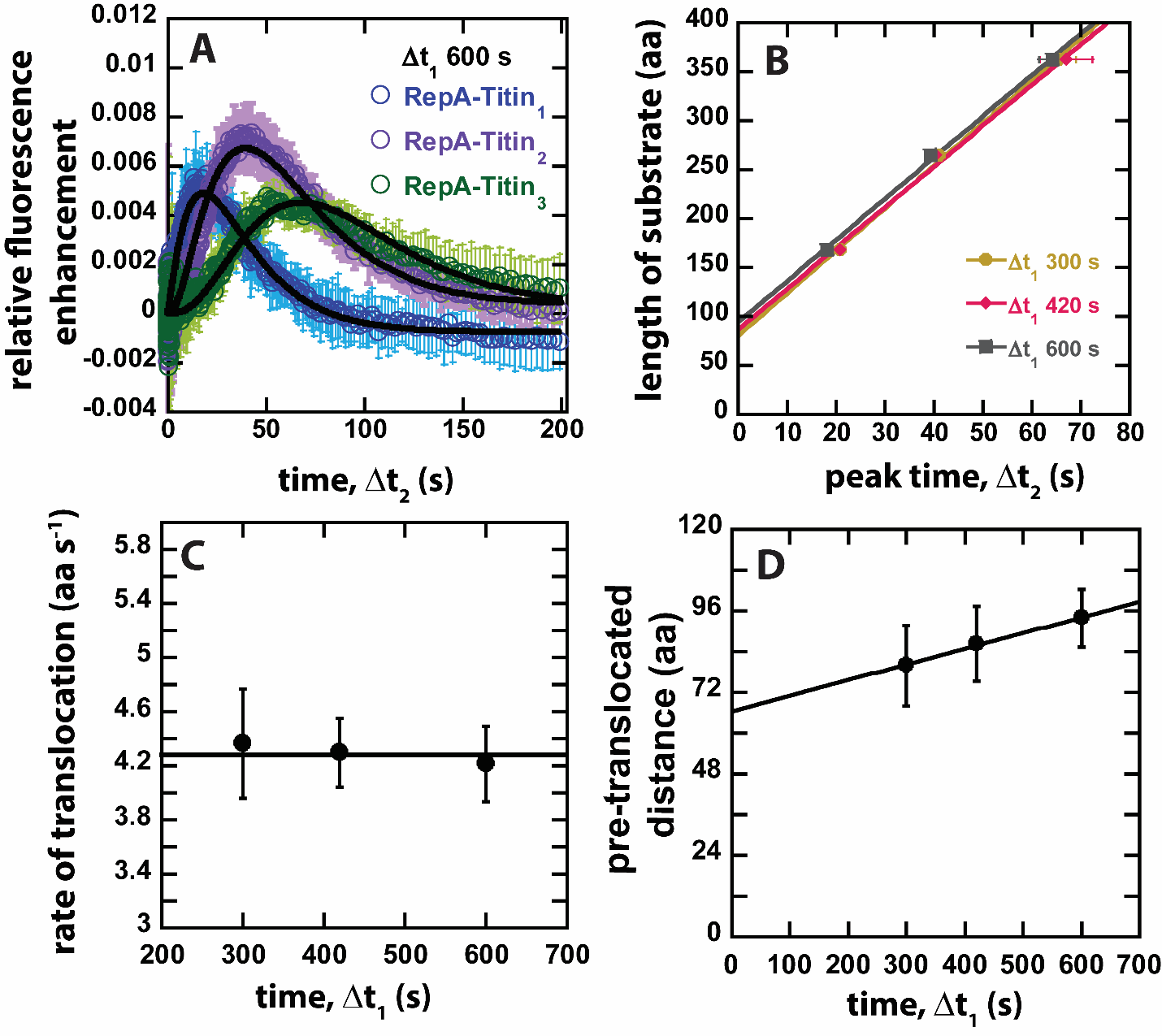

Supplemental Figure 7: ClpB catalyzed protein unfolding at approx. 3:1 ATP:ATPγS. Sequential mixing stopped-flow experiments were carried out as described in Fig. 3 A with the exception that Syringe 2 contains 600 μM ATPγS. Thus, after both mixing events, the final concentrations are 2 μM ClpB, 50 nM RepA-Titin_X_, 150 μM ATPγS, 500 μM ATP, and 20 μM α-casein. A) Representative time-courses from the average of five or more sequentially collected time-courses for RepA-Titin_1_(blue), RepA-Titin_2_(purple), and RepA-Titin_3_(green) at Δt_1_ = 600 s. The black solid lines represent the best-fit line from fitting to Scheme 1. The fitting parameters obtained are *k_U_* = (0.055 ± 0.005) s^-1^ and *m* = (58.4 ± 3.8) aa. B) Total length of substrate as a function of peak time determined for Δt_1_ = 300, 420, and 600 s as described in Materials and Methods. Solid lines represent weighted linear fits yielding C) slope vs. Δt_1_ with solid line representing the weighted average of (4.3 ± 0.2) aa s^-1^ and D) the intercept vs. Δt_1_ fit to a linear equation with a slope = (0.05 ± 0.05) aa s^-1^ and intercept of (67 ± 23) aa. All data points and error bars represent the average and standard deviation determined from three replicates.

### Supplemental Figure 8.

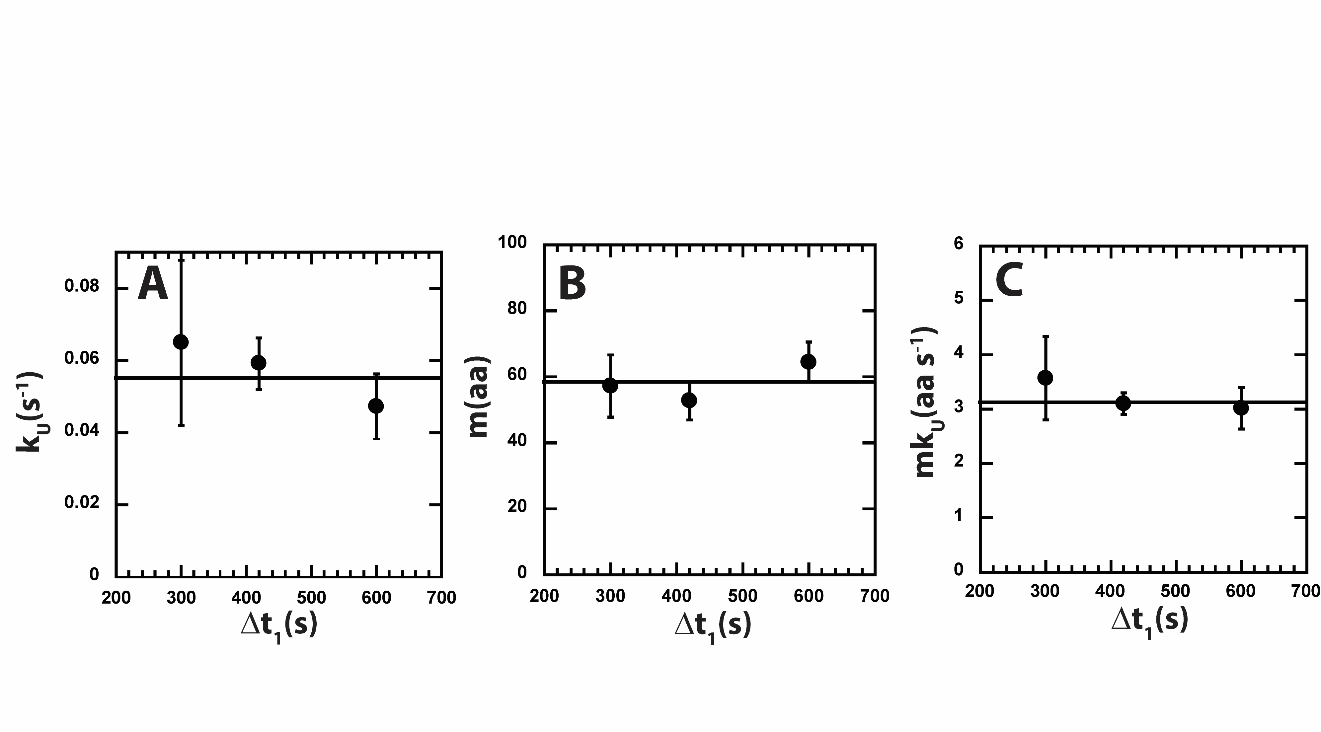

Supplemental Figure 8: Parameters from fitting to Scheme 1 for approx. 3:1 ATP: ATPγS. A) *k_U_* B) *m* C) *mk_U_* obtained from fitting to scheme 1 at each Δt_1_ are shown in solid black circles. The black solid line represents the weighted average value of *k_U_* = (0.055 ± 0.005) s^-1^, *m* = (58.4 ± 3.8) aa, and *mk_U_* = (3.1 ± 0.2) aa s^-1^.

**Confidence for RepA-Titin_1_**

In the predicted RepA-Titin_1_ structure, the most residues in the predicted α-helices at 34-40 and 59-72 have 90>pLDDT>70. Most residues in the β-sandwich at 79-161 have pLDDT>90.

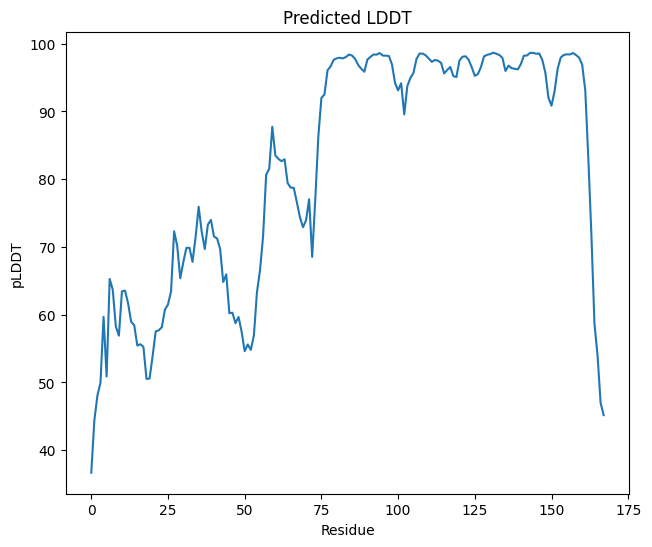

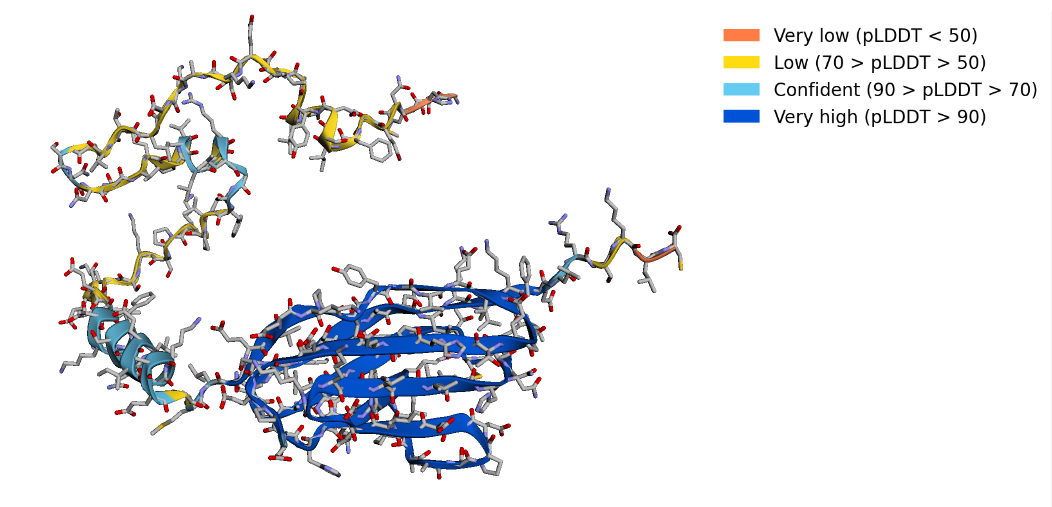

**Confidence for RepA-Titin_2_**

In the predicted RepA-Titin_2_ structure, most residues in the predicted α-helices at 34-40 and 60-72 have pLDDT<70. Most residues in the β-sandwiches at 79-161, 176-258 have pLDDT>90.

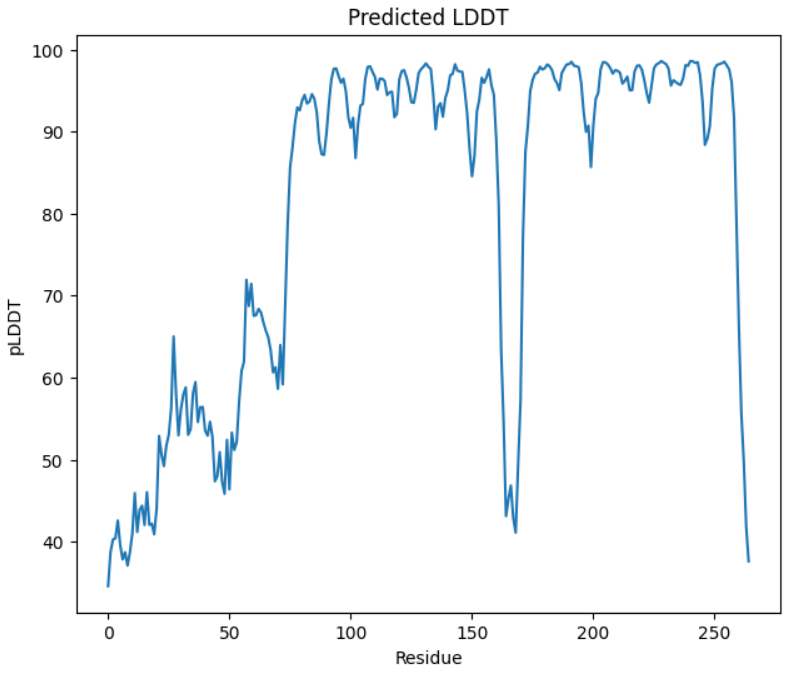

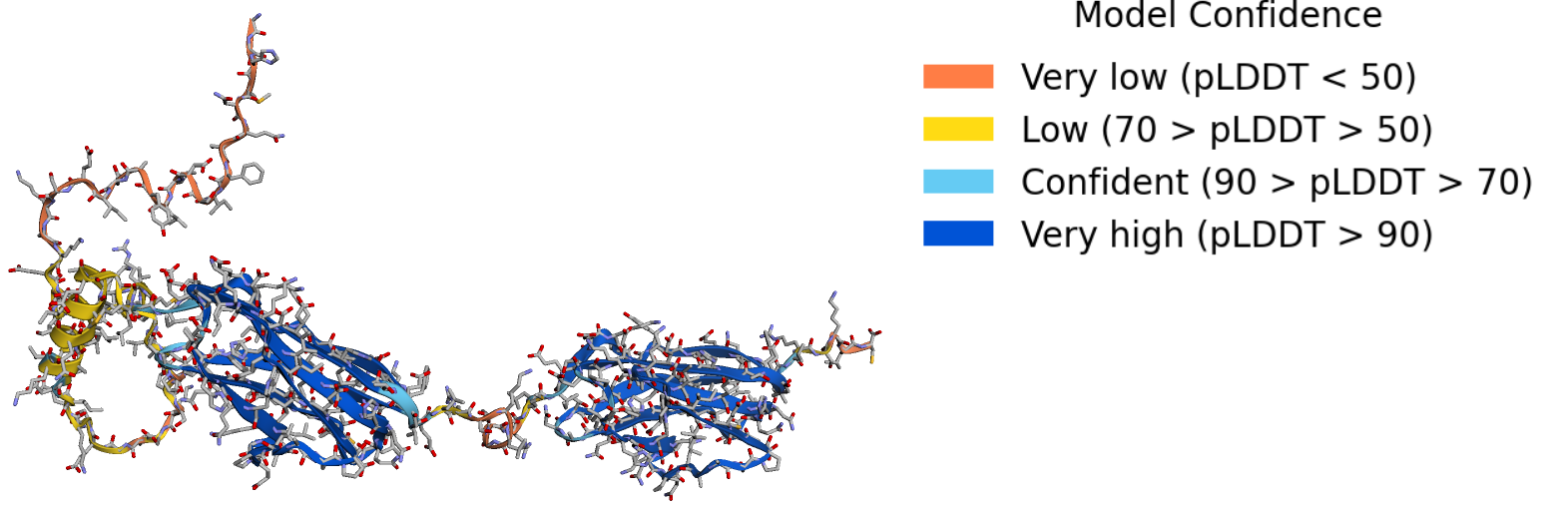

**Confidence for RepA-Titin_3_**

In the predicted RepA-Titin_3_ structure, the residues in the predicted α-helices at 34-40 and 60-72 have pLDDT<70. The vast majority of residues in the Most residues in the β-sandwiches at 79-161, 176-258, 273-355 have pLDDT>90.

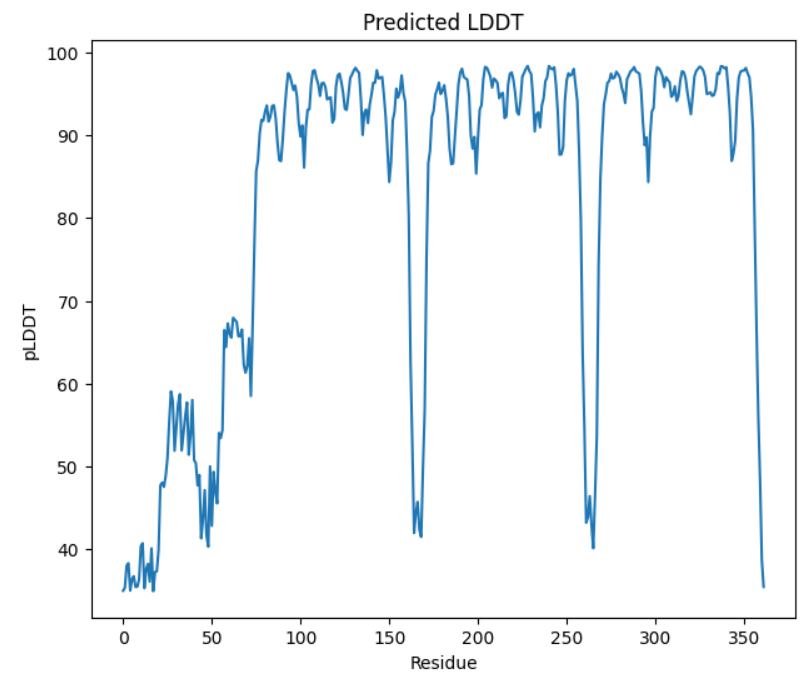

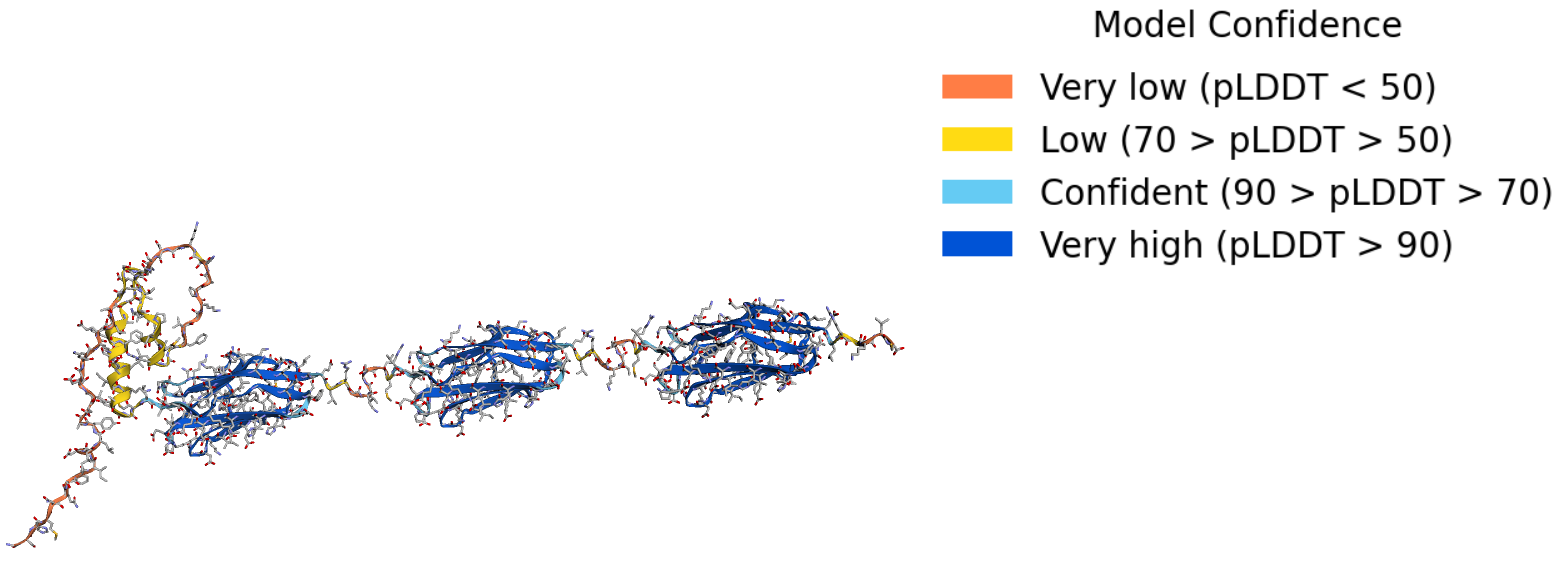

| **Substrate** | **Sequence Range** | **Secondary Structure** | **pLDDT for Most Residues** | **Model Confidence** |
| --- | --- | --- | --- | --- |
| RepA-Titin_1_ | 34-40 | α-helix | 70-90 | Confident |
|  | 59-72 | α-helix | 70-90 | Confident |
|  | 79-161 | β-sandwich | >90 | Very High |
|  | All other ranges | Any | <70 | Low or Very Low |
| RepA-Titin_2_ | 34-40 | α-helix | <70 | Low or Very Low |
|  | 60-72 | α-helix | <70 | Low or Very Low |
|  | 79-161 | β-sandwich | >90 | Very High |
|  | 176-258 | β-sandwich | >90 | Very High |
|  | All other ranges | Any | <70 | Low or Very Low |
| RepA-Titin_3_ | 34-40 | α-helix | <70 | Low or Very Low |
|  | 60-72 | α-helix | <70 | Low or Very Low |
|  | 79-161 | β-sandwich | >90 | Very High |
|  | 176-258 | β-sandwich | >90 | Very High |
|  | 273-355 | β-sandwich | >90 | Very High |
|  | All other ranges | Any | <70 | Low - Very Low |
